# Metabolic syndrome in New Zealand Obese mice promotes microglial-vascular interactions and reduces microglial plasticity

**DOI:** 10.1101/2023.10.04.560877

**Authors:** Michael MacLean, Olivia J. Marola, Travis Cossette, Cory Diemler, Amanda A. Hewes, Kelly J. Keezer, Kristen D. Onos, Gareth R. Howell

## Abstract

Metabolic syndrome (MetS) puts patients more at risk for neurodegenerative diseases such as Alzheimer’s disease (AD). Microglia are implicated as causal factors in AD, however, the effect of MetS on microglia has not been characterized. To address this, we contrasted New Zealand Obese (NZO) with C57BL/6J (B6J) mice in combination with a high fat/high sugar diet (HFD). Irrespective of diet, NZO mice displayed a broader array of MetS-relevant phenotypes compared to B6J mice fed a HFD. Single cell RNA-sequencing of microglia predicted transcriptional shifts indicative of reduced responsiveness and increased vascular interactions in NZO, but not B6J HFD mice. Significant cerebrovascular fibrin deposition and increased perivascular accumulation of microglia were observed in NZO relative to B6J HFD mice. Further, compared to the widely used B6J.*APP/PS1* mice, NZO.*APP/PS1* exhibited increased amyloid plaque sizes alongside an increase in microhemorrhages. Overall, our work supports a model whereby MetS alters microglia-vascular interactions, compromising microglial plasticity.

**Introduction:** Metabolic syndrome (MetS) is a combination of three or more metabolic impairments such as dyslipidemia, hyperglycemia, hypertension, and increased waist circumference (1). Patients displaying aspects of MetS are more at risk for neurodegenerative disorders including Alzheimer’s disease (AD) (2-6). Central to this risk may be the influence of MetS and Type 2 Diabetes (T2D) on peripheral and central immune cell states and function (7,8).

Burgeoning evidence has implicated critical roles for microglia, central nervous system (CNS) resident macrophages, in neurodegeneration (9-15). Microglia play roles in pathogen surveillance, debris phagocytosis, synapse regulation, and recently have been shown to support the cerebral vasculature (16-21). Current efforts have uncovered remarkable heterogeneity of microglial responses through single cell RNA-sequencing (scRNA-seq) (9,11,15,22). Increased disease-associated microglia (DAM) (11,12) and interferon response microglia (IRM) states (23) have been documented in neurodegeneration. The abundance of these states is heavily dependent on additional factors including age, sex, and genetic context (15,22-25). However, the effect of MetS on microglial states has yet to be determined.

To address this, we investigated MetS-induced changes in microglial transcriptional states using scRNA-seq. We utilized combinations of genetic and environmental models relevant to MetS contrasting New Zealand Obese (NZO/HlLtJ) mice, a polygenic model of obesity (26-28), with the commonly used C57BL/6J (B6J) mice fed a high fat/high sugar diet (HFD). Microglia scRNA-seq suggested MetS resulted in reduced microglial responsiveness in NZO mice relating to vascular interactions and function. In support of this, NZO but not B6J mice exhibited fibrin deposition within the cerebrovasculature. NZO mice also showed reduced microglia responses to an acute lipopolysaccharide (LPS) challenge compared to B6J mice, and larger amyloid-β plaques and increased microhemorrhages in the presence of *APP/PS1* transgenes (NZO.*APP/PS1*) compared to B6J.*APP/PS1* mice. In summary, MetS appeared to impair microglial plasticity, which could potentially drive neurodegenerative disease pathology.

**Results:** *Metabolic syndrome (MetS) caused subtle changes in abundances of microglia states.:* To determine how MetS affects microglia, we fed NZO and B6J mice a HFD or a standard diet (SD) from 2-9 months of age (mo). NZO mice showed signs of MetS at 2mo (Figure 1A-E). At 9mo, NZO mice displayed increased age-and diet-associated metabolic impairments relative to B6J mice, including weight gain, dyslipidemia, high blood pressure, and hyperglycemia (Figure 1F-J). These data indicate that the NZO strain better models complex endophenotypes observed in humans with MetS compared to HFD-fed B6J mice. Figure 1.
Strain-dependent effects of both high-fat diet and aging on characteristics of metabolic syndrome. **a.** Body weight at 2mo. **b.** Fasted cholesterol at 2mo. **c.** Fasted triglycerides at 2mo. **d.** Hemoglobin A1c (HbA1c) at 2mo. **e.** Blood pressure at 2mo. **f.** Body weight of mice fed either SD or HFD from 2-9mo. **g.** Fasted cholesterol at 9mo. **h.** Fasted triglycerides at 9mo. **i.** HbA1c at 9mo. **j.** Blood pressure at 8mo. In **a**–**e**: two-way ANOVA with post-hoc Tukey’s test. In **a,** *N*=16 NZO (7M,9F), *N*=23 B6J (11M,12F) mice. In **b-c**, *N*=7 NZO (4M,3F), *N*=8 B6J (4M,4F) mice. In **d**, *N*=8 NZO (4/sex), *N*=9 B6J (4M,5F) mice. In **e**, *N*=4M mice/strain. In **f**, mixed effects model with repeated measures. *N*=7M NZO (4SD,3HFD), *N*=9F NZO (4SD,5HFD), *N*=11M B6J (5SD,6HFD), *N*=12F B6J (7SD,5HFD). In **g-h**, *N*=7M NZO (4SD,3HFD), *N*=9F NZO (4SD,5HFD), *N*=10M B6J (5SD,5HFD), *N*=12F B6J (7SD,5HFD). Dashed lines and represent values > 200mg/dL. In **i**, *N*=7M NZO (4SD,3HFD), *N*=9F NZO (4SD,5HFD), *N*=10M B6J (4SD,6HFD), *N*=11F B6J (6SD,5HFD). Red line indicates diabetic HbA1c, gray line indicates prediabetes. In **j**, *N*=13M NZO, *N*=9M B6J mice. All data shown are Mean*±*SEM.

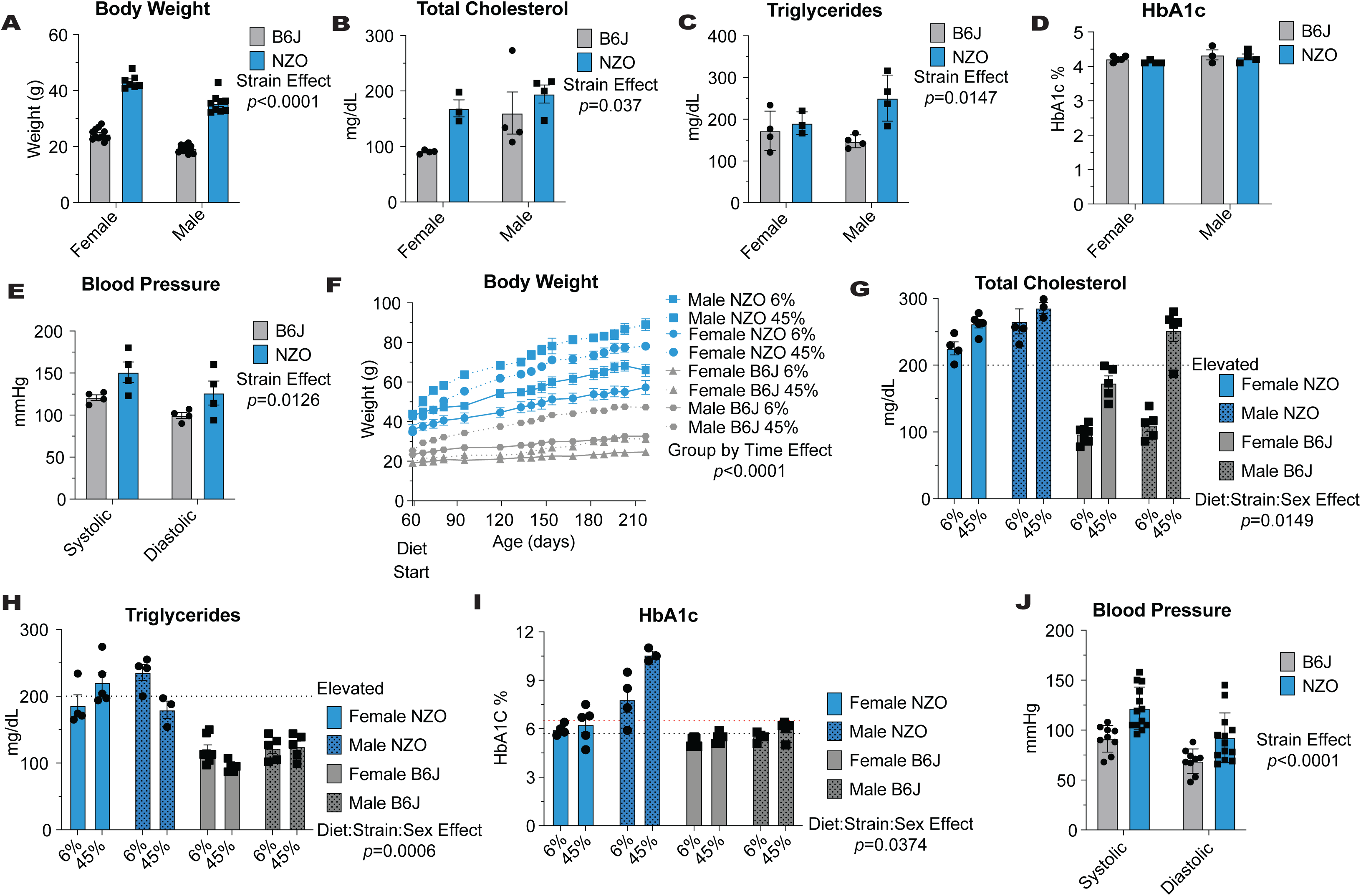 To determine the effects of strain (NZO, B6J) and diet (SD, HFD) on myeloid cell transcriptional states, we performed scRNA-seq of CD11B^+^ cells (24,29) from brains of 2 and 9mo male and female B6J, B6J HFD, NZO, and NZO HFD mice (Figure 2A). Microglia represented ∼75% of all captured CD11B^+^ cells in B6J mice, and ∼88% in NZO mice, irrespective of diet (Figure S1A-D) (30,31). The remaining cells consisted mainly of monocytes, macrophages, NK cells, and neutrophils (Figure S1D). Re-clustering only microglia resulted in 18 clusters (Figure S1E-F) with further annotation as: homeostatic (H, clusters: 0-6,9, 13,14, and 16), proliferating (cluster 17), *Hexb* high (HexB, cluster 10), disease-associated (DAM, cluster 8), *Ccl4 Ccl3* high DAM (*Ccl4*^+^ *Ccl3*^+^ DAM, cluster 7), major histocompatibility enriched (MHC, cluster 16), interferon responsive microglia (IRM, cluster 12) and *Klf2 Tcim* high (*Klf2^+^ Tcim^+^,* cluster 13) (Figure 2B-D, Figure S1E-F) (11,12,22,24). In contrast to amyloid-(11,12,24) and aging-related (22) studies, MetS (NZO strain and/or HFD) did not cause significant changes in the percentages of DAM, IRM or MHC clusters (Figures 2E, S2A). However, the abundance of DAM, MHC, and *Klf2*^+^ microglia changed with age, regardless of strain or diet (Figure 2E, Figure S2A-B). Figure 2.
Profiling myeloid transcriptional states in a cohort of varying metabolic impairment. **a.** Experimental design scheme; created with BioRender.com. **b.** Dimensionality reduction plot (UMAP) of microglia colored by cluster. **c.** UMAPs of microglia colored by SCT-normalized expression of marker genes. **d.** Dot plot of marker genes associated with each annotated state. **e.** UMAP of microglia colored by annotated state. In **a-e**, 9mo mice: *N*=3M NZO/diet, *N*=4F NZO/diet, *N*=4M B6J/diet, *N*=3F B6J/diet; 2mo mice: *N*=3 NZO/sex, *N=*7 B6J (3M,4F).

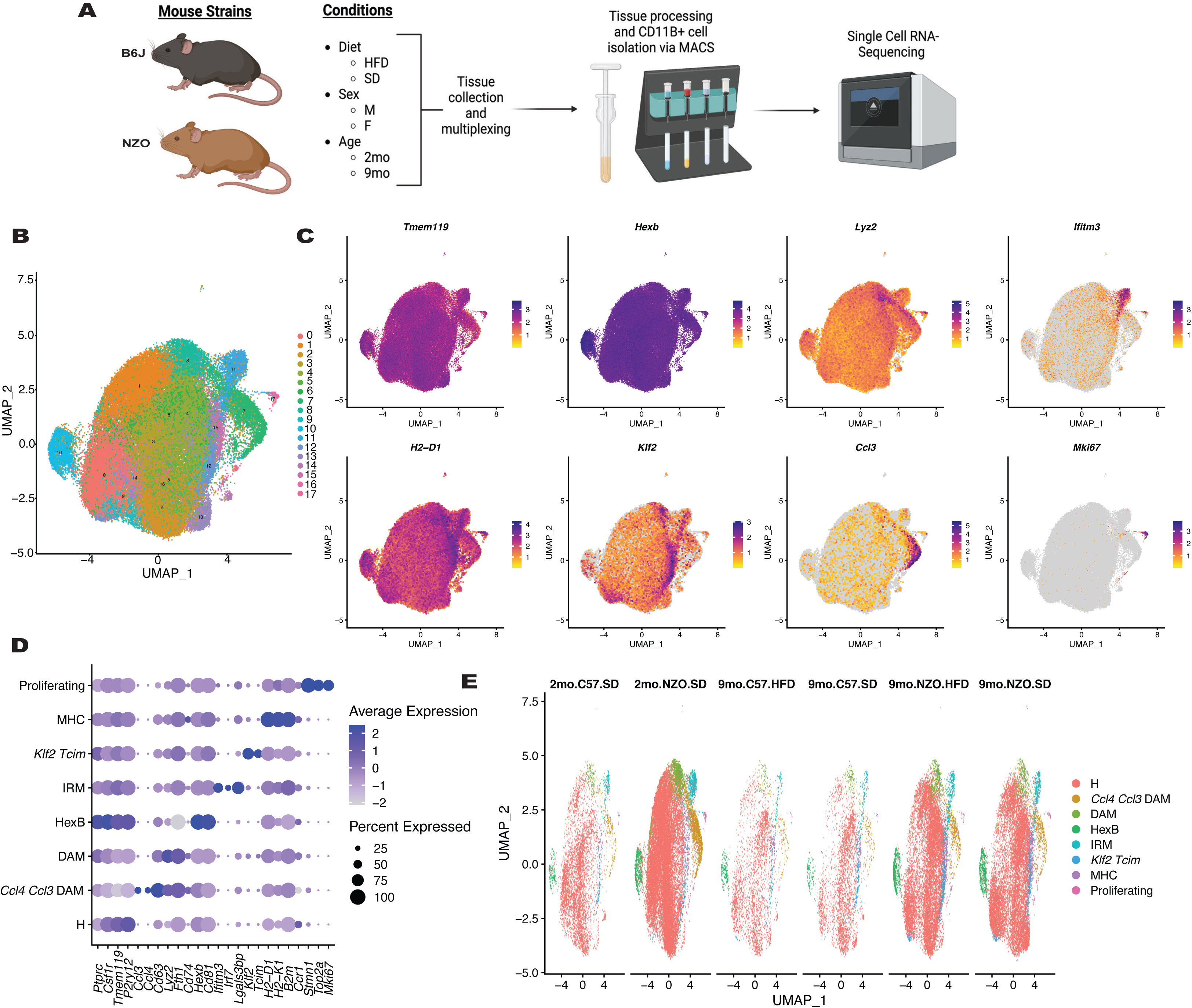

*High fat diet altered stress response and cell-cell communication gene expression in NZO, but not B6J microglia:* Despite the lack of MetS-induced shifts in microglial states (Figure 2E), we reasoned that MetS may alter gene expression across all microglia. To determine this, we utilized a pseudobulking strategy followed by differential expression analyses, separately assessing the strain (NZOvB6J), and diet (HFDvSD) or aging (9v2mo) effect within each strain (Figure S3A-B)(32,33). First, we determined the HFD effect across both NZO and B6J mice and found that there were 141 differentially expressed genes (DEGs) in all microglia, which were enriched in gene sets associated with cell viability, migration, and proliferation (Figures 3A, S4A-C). However, when HFD effects were analyzed separately for each strain, B6J microglia did not show a substantial response to chronic HFD exhibiting only 2 DEGs in comparison with >300 DEGs in NZO microglia (Figure 3A,S4D). These NZO DEGs were associated with endothelial and immune cell signaling, and heat shock stress (Figure 3B-D). Together, these data suggest HFD caused a significant stress response in microglia in NZO, but not B6J mice. Figure 3.
HFD promotes stress-responses and alters cellular communication pathways in NZO but not B6J microglia. **a.** Bar chart summarizing the number of DEGs associated with HFDvSD across all microglia, within NZO microglia alone, and within B6J microglia alone. **b.** IPA graphical summary of NZO HFDvSD microglia DEGs. **c.** Violin plots of selected HFDvSD NZO DEGs. **d.** Enrichment GO term plot for NZO HFDvSD microglia DEGs. In **a-d**, *N*=3M NZO/diet, *N*=4F NZO/diet, *N*=4M B6J/diet, *N*=3F B6J/diet mice.

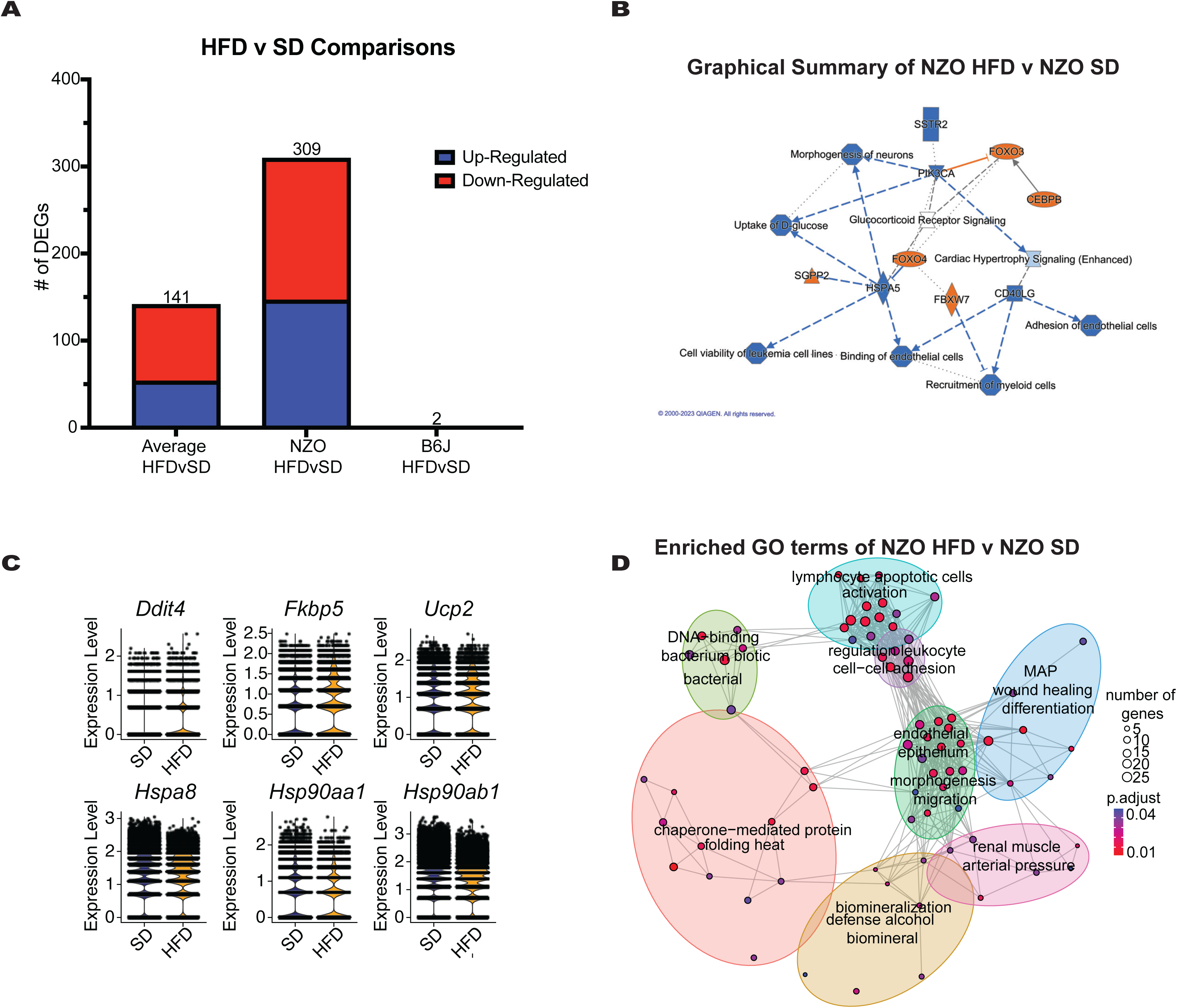

*Aging differentially effected NZO and B6J microglia:* To assess transcriptional programs in NZO compared to B6J, we first identified DEGs comparing NZO to B6J microglia at 2 or 9mo. DEGs identified at both ages included genes that have been implicated in regulating microglial and macrophage responses, such as *Angptl7*, *Itgam,* and *Fcrls* (Figure 4A-B,S5A-D) (22,34-36). Interestingly, *Apoe*, genetic variations in *APOE* increase risk for AD (37,38), was significantly greater in NZO compared to B6J microglia at both ages (Figure 4C). At 9mo, DEGs were associated with immune responses, vascular interactions, and cell migration (Figure 4D-E), suggesting these processes are perturbed in NZO but not B6 microglia. When microglia states were analyzed separately, differentially expressed genes detected in all microglia were primarily driven by homeostatic microglia, downsampling indicated this was independent of the number of microglia within each state (Figure S5A-B). Figure 4.
NZO and B6J microglia exhibit strain- and age- associated transcriptional differences. **a.** Venn diagram displaying overlap of NZOvB6J DEGs at 2 and 9mo. **b.** Violin plots of a subset of NZOvB6J 9mo DEGs. **c.** UMAP plots for NZO and B6J microglia. Colored by SCT-normalized *Apoe* expression. **d.** Enrichment GO term plot of 9mo NZOvB6J microglia DEGs. **e.** IPA graphical summary of 9mo NZOvB6J DEGs. **f.** Venn diagram of 9v2mo DEGs identified in NZO and B6J microglia. **g.** Violin plots of selected 9v2mo NZO microglia DEGs. Enrichment GO term plots for 9v2mo B6J (**h**) or NZO (**i**) microglia DEGs. 9mo mice: *N*=7 NZO (3M,4F), *N*=7 B6J (4M,3F). 2mo mice: *N*=6 NZO mice (3M,3F), *N=7* (3M,4F) B6J mice.

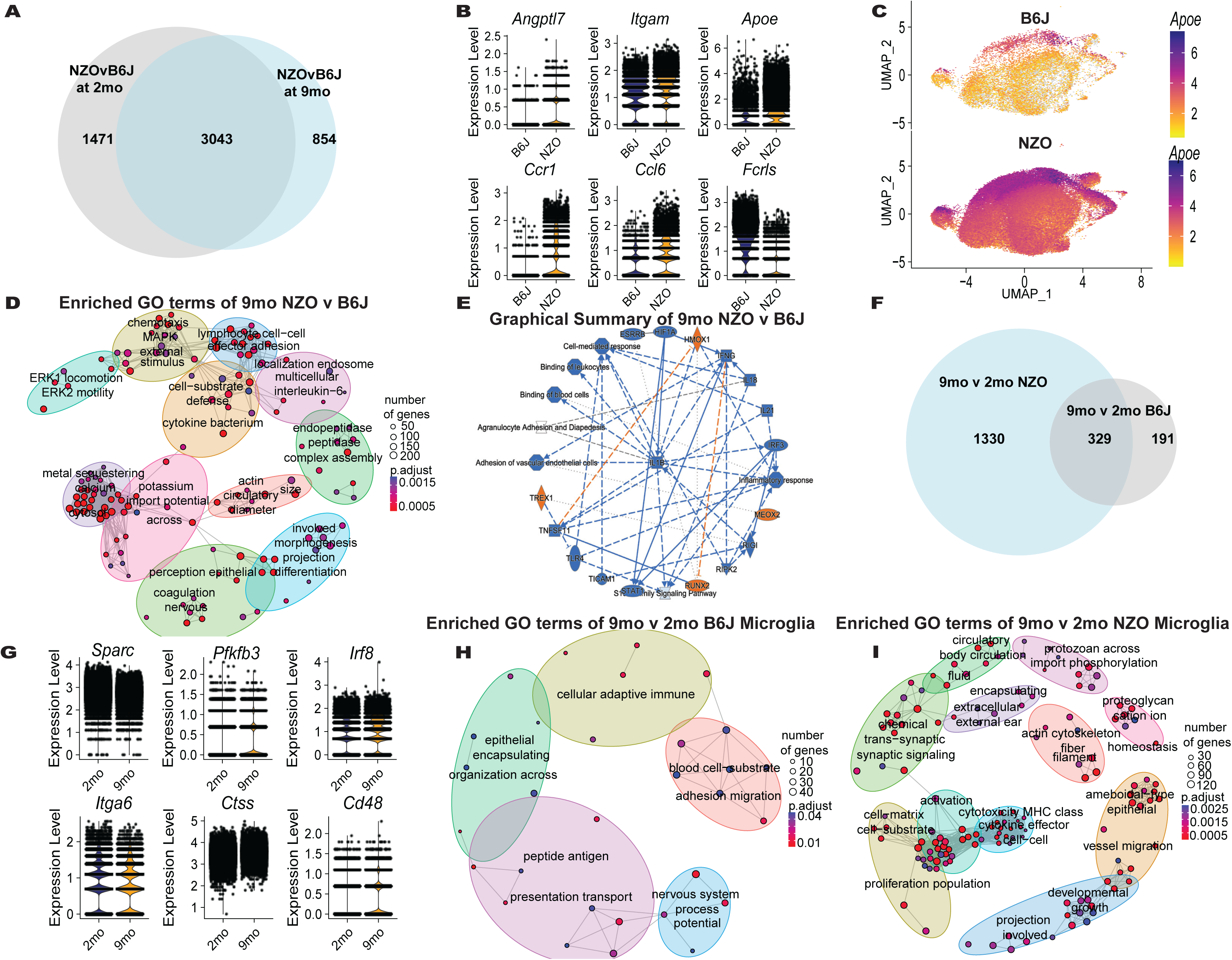 Next, to better understand how transcriptional programs were influenced by aging, we compared 9 to 2mo microglia from either NZO or B6J mice (Figure 4F). Most aging related transcriptional changes were again detected within the homeostatic microglia state when analyzed separately (Figure S6A-B). Both strains showed aging-associated changes in *Itga6*, *Ctss*, *Cd48*, and antigen processing and presentation pathways (Figure 4F-I). However, NZO microglia exhibited more aging-associated DEGs than B6J including *Sparc*, *Pfkfb3,* and *Irf8*; which have been implicated in regulating synaptic function, glycolysis, and microglial identity respectively (Figure 4F- G) (39-41). In addition, NZO microglia exhibited age-dependent changes in cytotoxicity, wound healing, and circulation pathways (Figure 4I). Interestingly, Ingenuity Pathway Analysis (IPA) of upstream regulators predicted aging-associated upregulation of interferon signaling regulators in B6J microglia, but regulators associated with environmental stress and tissue repair in NZO microglia (Figure S6C-E).

*NZO microglia displayed increased association with blood vessels:* The expression of genes regulating myeloid-endothelial interactions including *Itgam*, *Ccr1, P2ry12*, and *Ccr5* (16,35,36,42), was higher in NZO relative to B6J microglia (Figure 4). To probe this further, we performed immunohistochemistry to localize microglia and vasculature within the cortex and hippocampus of 9mo NZO and B6J mice fed SD or HFD. We found that while the numbers of TMEM119^+^DAPI^+^ microglia in the hippocampus or cortex did not differ across strains or diets, the percentage of CD31^+^ area covered by microglia was significantly higher in NZO relative to B6J mice (Figure 5A-E). HFD did not modulate this phenomenon (Figure 5B-E). scRNA-seq analyses predicted fibrin(ogen) to be an upstream regulator of aging-associated DEGs in NZO microglia (Figure 5F). Fibrin is absent in the healthy CNS but can deposit in the perivascular space and within vessels in conditions of stress (38,43,44). We found NZO vessels exhibited peri-vascular and vascular deposition of fibrin in the hippocampus and cortex, while B6J vessels did not (Figure 5G-I). Furthermore, many of these fibrin^+^ vessels had microglia juxtaposed (Figure 5G). Altogether, these data suggest that vessel stress signals may promote microglia-vascular interactions in NZO mice. Figure 5.
NZO vasculature displays fibrin deposition concomitant with increased coverage by microglia. **a.** Representative images of TMEM119 and CD31 staining in the hippocampus. Quantification of microglia in the hippocampus (**b)** and cortex (**d**). Quantification of the percentage of CD31 colocalized with TMEM119 in the hippocampus (**c**) and cortex (**e**). **f.** IPA upstream regulator analysis of 9v2mo NZO microglia DEGs. **g.** Representative images of TMEM119, CD31, and fibrin. Insets are high resolution confocal imaging of noted area. Quantification of the percentage of CD31 colocalized with fibrin in the hippocampus (**h**) and cortex (**i**). *N*=8 NZO (2/sex/diet) and *N*=8 B6J (2/sex/diet) mice. In **b**-**c**, independent two sample two-sided *t* test. In **e**, **h-i**, Mann-Whitney test. SD: closed circles, HFD: open circles. Data are presented as Mean±SEM.

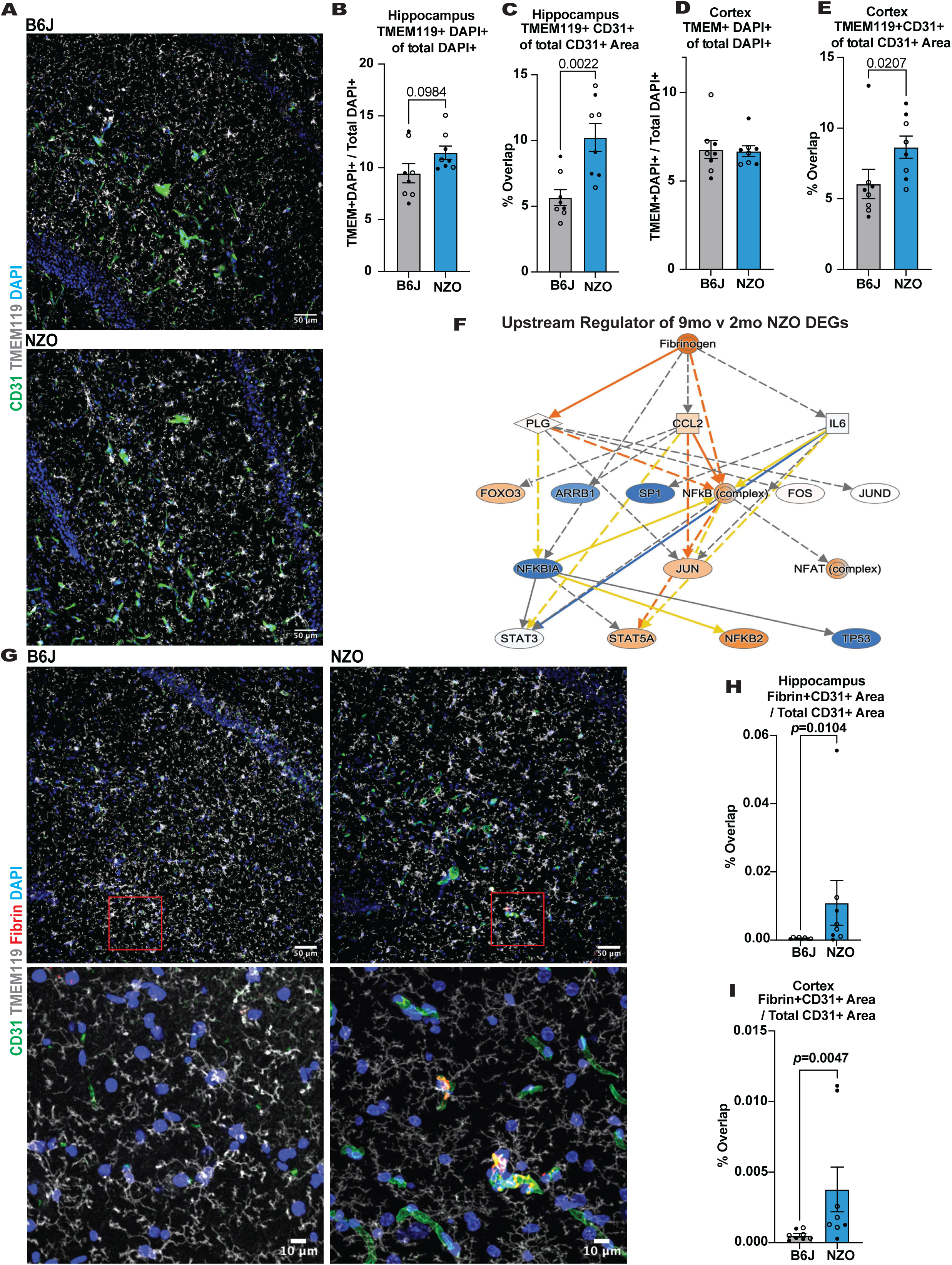

*NZO mice displayed a dampened response to LPS:* As NZO microglia appeared to have a reduction in DEGs relevant to immune responses compared to B6J microglia (Figure 4,S5), we sought to probe the responsiveness of NZO microglia to an acute inflammatory challenge. LPS was administered to 4mo NZO and B6J mice, and bulk hemibrain RNA-seq (Figure 6A) was performed. As we suspected from the observed fibrin deposition, NZO animals exhibited altered CNS expression of vascular associated pathways and genes including *Edn1, Angpt1,* and *Serpine1* (Figure 6B-C) even with PBS-treatment. LPS-treated NZO animals also displayed fewer DEGs than LPS-treated B6J mice (Figure 6D). The strain-dependent LPS effects suggested potential differences in microglial responses, as NZO animals displayed no change in *Cx3cr1* expression with LPS treatment (Figure 6E). Furthermore, IPA upstream regulator and pathway analyses predicted strain-dependent differences in the LPS induction of genes involving macrophage recruitment, antigen processing and presentation, and aggregation of cells (Figure 6F). These data indicate that compared to B6J, NZO mice display altered microglial responses to not only HFD, but also to an acute insult such as LPS. Figure 6.
NZO animals display increased expression of hemibrain vascular associated genes and an altered central nervous system response to an acute LPS treatment. **a.** Experimental schematic of CNS responsiveness to LPS. Created with BioRender.com. **b.** Bar chart of DEGs associated with each comparison. **c.** Top IPA regulatory effect for NZOvB6J DEGs. **d.** Venn diagram displaying DEGs associated with the LPS response in each strain. **e.** Volcano plot of the Strain-by-Treatment interaction effect. Top 10 genes by significance are labeled. Genes are colored by significance. **f.** IPA graphical summary of the Strain:LPS interaction effect. In **a-f**, N=3F mice/strain/treatment.

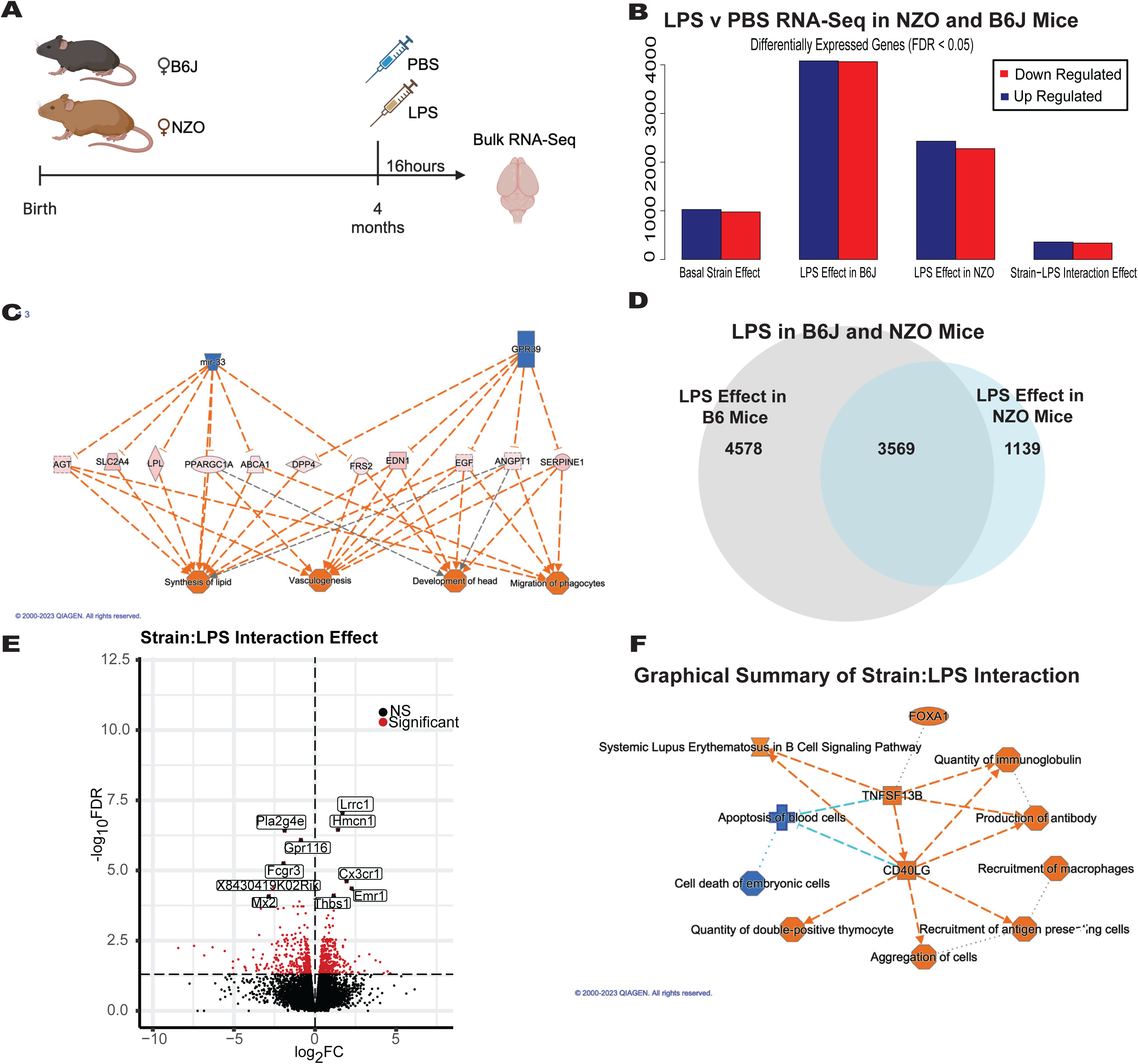

*NZO.APP/PS1 mice displayed larger amyloid plaques and increased incidences of microhemorrhages:* MetS has been demonstrated to increase risk for AD and related dementias (2,4,5,45). We hypothesized that this may be due to reduced responsiveness of microglia to AD-relevant insults such as amyloid deposition. To test this, we backcrossed the commonly used amyloid-inducing transgenes (*APP/PS1*) (46) from B6J to NZO resulting in ∼98.375% congenicity. Unexpectedly, male NZO.*APP/PS1* mice exhibited pronounced weight loss, likely associated with exacerbation of T2D (Figure S7A-E). To avoid this confound, we primarily focused our analyses on amyloid-related microglia responses in female mice. At 8mo, in comparison to B6J.*APP/PS1* mice, female NZO.*APP/PS1* mice exhibited fewer, but larger, amyloid plaques in both the hippocampus and cortex, which were positive for the dystrophic neurite marker LAMP1 (Figure 7A-L). Similar results were observed in several male NZO.*APP/PS1* mice that survived to 8mo (Figure S7F-G). There was a small but significant increase in total IBA1^+^ area in the hippocampus, but not the cortex of NZO.*APP/PS1* compared to B6J.*APP/PS1* mice (Figure 7F,K). However, the area of plaque covered by microglia were the same in both regions (Figure 7G, L). Figure 7.
NZO mice exhibit fewer, but larger neuritic amyloid plaques, without differences in microglia coverage of plaques. Representative images of 6E10 amyloid-β staining with IBA1 (**a**) or regions of interest with LAMP1 co-staining (**b**) in the hippocampus. **c.** Quantification of 6E10^+^ area in the hippocampus (**c**) and cortex (**h**). Quantification of 6E10^+^ counts in the hippocampus (**d**) and cortex (**i**). Quantification of the average size of 6E10^+^ objects in the hippocampus (**e**) and cortex (**j**). Quantification of IBA1^+^ area in the hippocampus (**f)** and cortex (**k**). Quantification of the percentage of 6E10^+^ area colocalized with IBA1^+^ area in the hippocampus (**g**) and cortex (**l**). **m**. Representative images of Prussian blue staining with higher magnification image of the inset. **n.** Quantification of Prussian blue^+^ microhemorrhages per brain section. SD: closed circles, HFD: open circles. In **a-l**, *N*=4F NZO*.APP/PS1*, *N*=3F B6J*.APP/PS1* mice. Independent two sample two-sided *t* test. In **m-n**, *N*=4/strain/diet WT mice and *N*=4F NZO, *N*=3F B6J *APP/PS1* mice. Two-way ANOVA with post-hoc Tukey’s test. Data are presented as Mean*±*SEM.

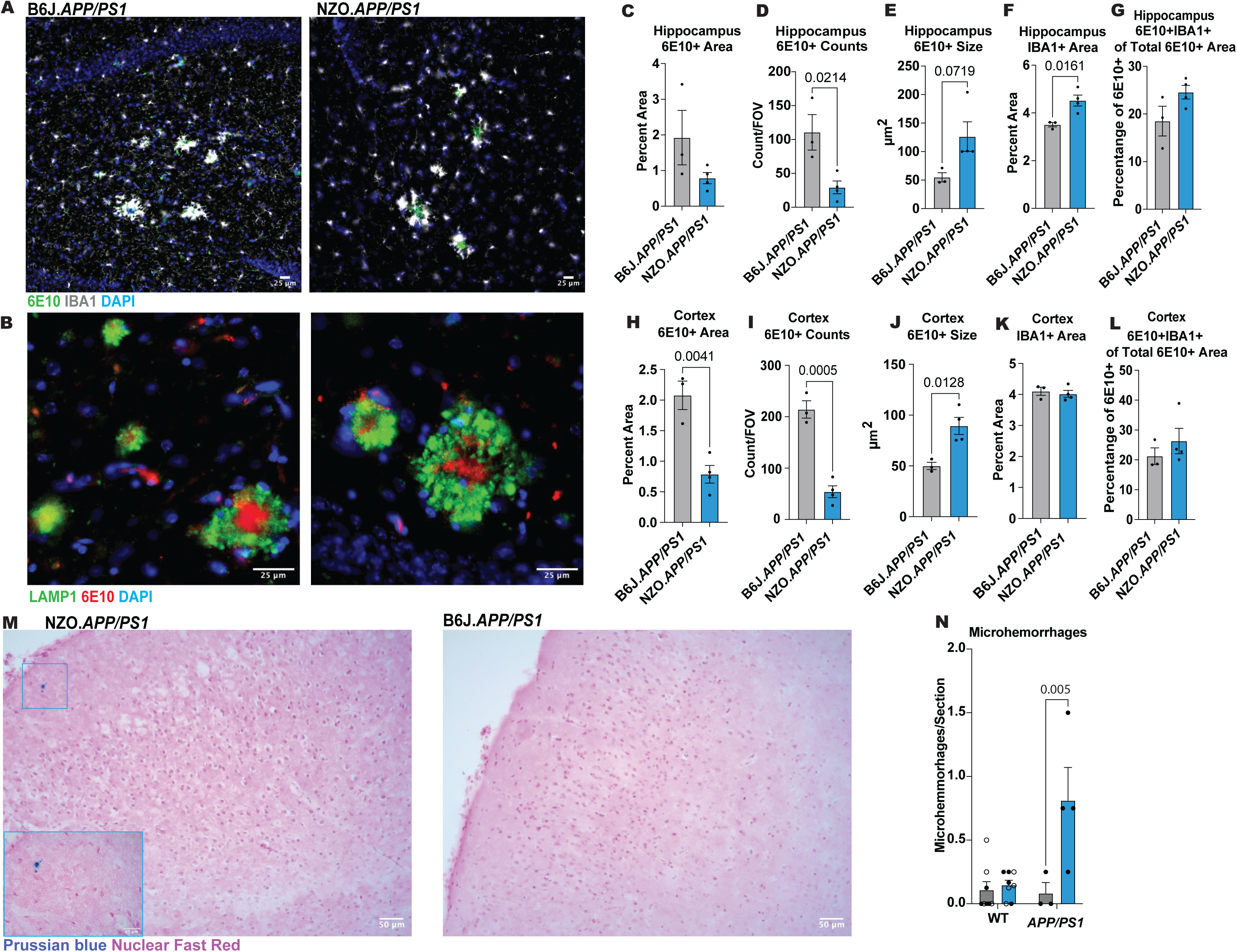 Previous studies have shown microglia depletion drives development of cerebral amyloid angiopathy (CAA) (14,47). We wondered whether the reduced responsiveness of NZO microglia would result in increased CAA. However, extensive CAA was not present in NZO.*APP/PS1* or B6J*.APP/PS1* mice (Figure 7A, S7F-G). Our previous findings of increased vascular stress in NZO (Figures 5-6) suggested blood-brain-barrier integrity may be more compromised in NZO*.APP/PS1* compared to B6J.*APP/PS1* mice. To assess this, we stained brain tissue with Prussian blue, which marks areas of iron deposition, and found that NZO*.APP/PS1* mice displayed increased incidence of microhemorrhages throughout the brain relative to either B6J.*APP/PS1* mice or their WT littermate controls (Figure 7M-N). Together, these data support that in a context of MetS, amyloid plaques are larger and the cerebrovasculature is more prone to microhemorrhages, outcomes which may increase risk for diseases such as AD.

**Discussion:** Our work focused on the relationship between MetS and alterations in microglial responses using mice exhibiting varying aspects of MetS (26-28). Consistent with previous reports, NZO mice displayed most aspects of MetS, and this was exacerbated with HFD. However, HFD-fed B6J mice exhibited dysfunctional metabolic measures associated only with pre-diabetes and obesity, identifying NZO mice as a more appropriate model of MetS than B6J HFD mice. We profiled 83,757 microglia and unexpectedly, MetS did not significantly shift the proportion of previously identified microglial states (11,12,22,24). We noted an increase in MHC and *Klf2^+^* transcriptional states and a decrease in both DAM populations between 2 and 9mo. This is consistent with previous reports of age-related alterations in MHC and DAM microglia populations (48).

We identified significant strain-specific changes in gene expression programs across all microglia, and within specific states. Strikingly, when we analyzed differential expression within each state, cells within the homeostatic clusters exhibited the greatest number of DEGs even after downsampling. This suggests that these transcriptional changes, while not sufficient to alter state abundances, are potentially altering microglial function. Strain-specific differences in microglia are likely driven by the MetS endophenotypes exhibited by NZO as early as 2mo. NZO microglia exhibited significantly higher expression of *Apoe* compared to B6J or B6J HFD microglia. ApoE has been linked to the transition to a DAM (or activated response microglia) state (12,23), yet, there was no difference in the abundance of DAM between NZO and B6J mice. One explanation for this paradox may be that the increase in *Apoe* expression is in response to dyslipidemia, as ApoE has known roles in lipid metabolism (37,49,50). Furthermore, it is possible that NZO microglia are unable to fully transition to a DAM state yet still acquire DAM-like characteristics such as high *Apoe* expression. In addition to the aging-and HFD-independent strain differences, there were also significant aging-and HFD-dependent strain differences between NZO and B6J microglia. For instance, aging influenced NZO microglia more than B6J microglia through higher numbers of DEGs and a broader range of impacted pathways, including wound healing and cytotoxicity. One recent study suggested that NZO mice display enhanced aging-associated changes within peripheral immune populations —suggesting that NZO were a model of accelerated aging (51) — and our data also support this possibility.

A prevailing signature of both aging-and strain-associated analyses implicated microglia differential interactions with the vasculature. Further exploration through IHC highlighted that regardless of diet, NZO mice exhibited more perivascular microglia than B6J mice. Upstream regulator analysis predicted fibrin may a significant culprit behind aging-associated changes in NZO microglia and fibrin deposition was detected in NZO brains. Fibrin has previously been shown to be neurotoxic (35,36). Fibrin upregulates *Hmox1* expression in microglia (36) and upstream regulator analysis predicted activation of HMOX1-dependent inflammatory response signaling pathways when comparing 9mo NZO and B6J microglia. Furthermore, MetS has been associated with increased CCL5, which can recruit immune cells via CCR1/CCR5 to vasculature (52-54). NZO mice exhibit peripheral vascular stress (55), and NZO microglia display higher expression of *Ccr1* and *Ccr5*.

Collectively, these data predict MetS may increase vascular stress and fibrin deposition in the CNS, resulting in recruitment of microglia to the cerebrovasculature. This primary endophenotype may than render microglia less responsive to a secondary insult. To test this, we first used an acute LPS treatment and performed bulk RNA-seq on hemibrains. We found increased expression of vascular associated genes in NZO hemibrains relative to B6J hemibrains including *Serpine1*, *Edn1* and *Angpt1* which mediate vascular stress and fibrin accumulation (56-58). Further, fewer DEGs were identified in LPS-treated NZO mice compared to LPS-treated B6J. This provides support for decreased responsiveness in NZO microglia. For example, *Cx3cr1* did not change in LPS-treated NZO mice. *Cx3cr1* expression has been shown to decrease in neurodegeneration (11,12). Interestingly, recently published data suggest aged microglia display a dampened response to LPS, supporting the concept that NZO mice may be a model of MetS-dependent accelerated aging (59).

Following acute stimuli, we turned to a more chronic and disease-relevant inflammatory stimulus, amyloid deposition (12,14,23,60). We found that NZO*.APP/PS1* mice displayed a significant increase in plaque size, independent of numbers of plaque-associated microglia. Increased plaque sizes may impact larger regions leading to increased likelihood of cognitive dysfunction. Patients displaying MetS have accelerated plaque deposition, and MetS blood biomarkers correlate with the rate of cognitive decline in patients with MCI and dementia (61,62). One possibility for the increased plaque size is in NZO.*APP/PS1* compared to B6J.*APP/PS1* mice is inefficient plaque compaction or clearance by NZO microglia - that would fit with the model of MetS-dependent reduced responsiveness. A second possibility may be altered activity of insulin degrading enzyme (IDE). In addition to insulin degradation, IDE also contributes to plaque degradation (63). Microglia-specific or brain-wide *Ide* expression was not changed between NZO and B6J mice, however, a MetS-dependent increase in insulin, requiring degrading by IDE in NZO mice, may result in reduced amyloid-β degradation.

Emerging work has implicated microglia in regulating blood flow and closure of injured vascular barriers (16,18,64). Therefore, given the vascular stress and changes to microglia-vascular interactions in NZO mice, we investigated whether NZO.*APP/PS1* mice were susceptible to microhemorrhages. NZO*.APP/PS1* mice presented with microhemorrhages, which were rare in B6J.*APP/PS1* mice or WT littermate controls. Microhemorrhages are more common in AD patients than in control groups and were previously detected in *Ob/Ob APP/PS1* mice (65-68). One possible mechanism driving the microhemorrhages in NZO.*APP/PS1* mice relates to fibrin deposition. Fibrin is stabilized by amyloid-β potentiating vascular damage and blood-brain-barrier breakdown (38,69). Recent evidence suggests the amount of fibrin coverage of vessels correlates to cerebral microbleeds in patients with CAA (70). Yet, we did not observe extensive CAA in NZO.*APP/PS1* mice. It is possible microhemorrhages in NZO.*APP/PS1* mice are a result of the inability of microglia to clear fibrin and/or amyloid-β efficiently, alongside potential impairments in wound healing and vascular repair pathways may predispose NZO mice to increased microhemorrhages. Diabetes worsens wound healing and vascular repair pathways in a variety of pathological conditions (71) and NZO microglia exhibit age-associated alterations in wound healing pathways underscoring this possibility. These data provide further evidence that MetS reduces or dampens the responsiveness of microglia.

In summary, we have found that MetS in NZO mice fundamentally altered microglia responses even when compared to microglia of HFD-fed B6J mice. NZO microglia displayed age-associated transcriptional changes concomitant with increased perivascular association. These changes were associated with abnormal responses to an acute LPS challenge and chronic amyloid pathology altogether suggesting MetS reduced microglial plasticity. Overall, this also supported the hypothesis of accelerated aging in NZO compared to B6J. This work provides the foundation to investigate the mechanisms by which MetS compromises microglial responses, leading to increased risk for neurodegenerative disease such as Alzheimer’s disease.

## Acknowledgments

We gratefully acknowledge the contribution of the Single Cell Biology Laboratory, Clinical Chemistry and Genomic Technologies Cores at The Jackson Laboratory (JAX) for expert assistance with this publication. We thank Dan Skelly from JAX Computational Sciences for advice on single-nucleotide polymorphisms based demultiplexing of scRNA-seq data. We thank Bansri Patel from the JAX Summer Student Program and Matthew Cox, a Mount Desert Island High School student intern, for preliminary investigations into NZO*.APP/PS1* animals.

This study was supported in part by National Institutes of Health National Institute of Aging T32G062409A (M.M.), JAX Scholars Program (M.M.); The Jackson Laboratory startup funds (G.R.H.), a BrightFocus Foundation award G2020254 (G.R.H.), and ongoing philanthropic support from the Diana Davis Spencer Foundation and other anonymous donors (G.R.H.). G.R.H holds the Diana Davis Spencer Foundation chair for glaucoma research. The funders had no role in study design, data collection, analysis, decision to publish, or preparation of the manuscript.

## Author contributions

M.M, K.D.O., and G.R.H. designed the study. M.M. developed experimental protocols, completed *in vivo* and IHC experiments and analyses, performed microglia isolations, and scRNA-seq bioinformatics analyses. O.J.M, C.D., T.C., K.D.O. and K.J.K, and A.A.H., all contributed to experiments. A.H., M.M., and K.J.K. generated and maintained mice for this study. M.M, K.D.O., and G.R.H. wrote the manuscript. All authors approved the final version.

## Declaration of interests

The authors declare no competing interests

---

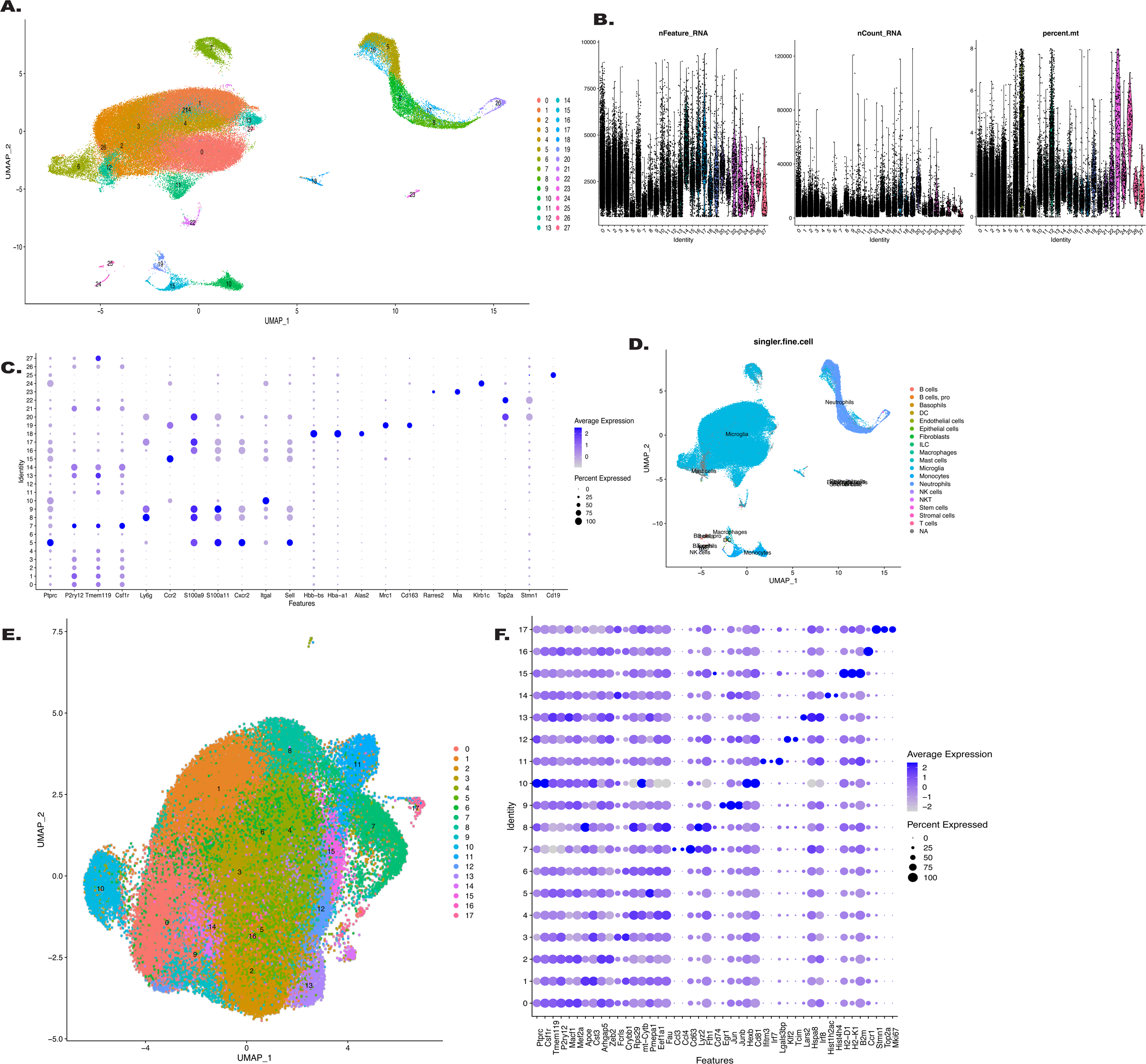

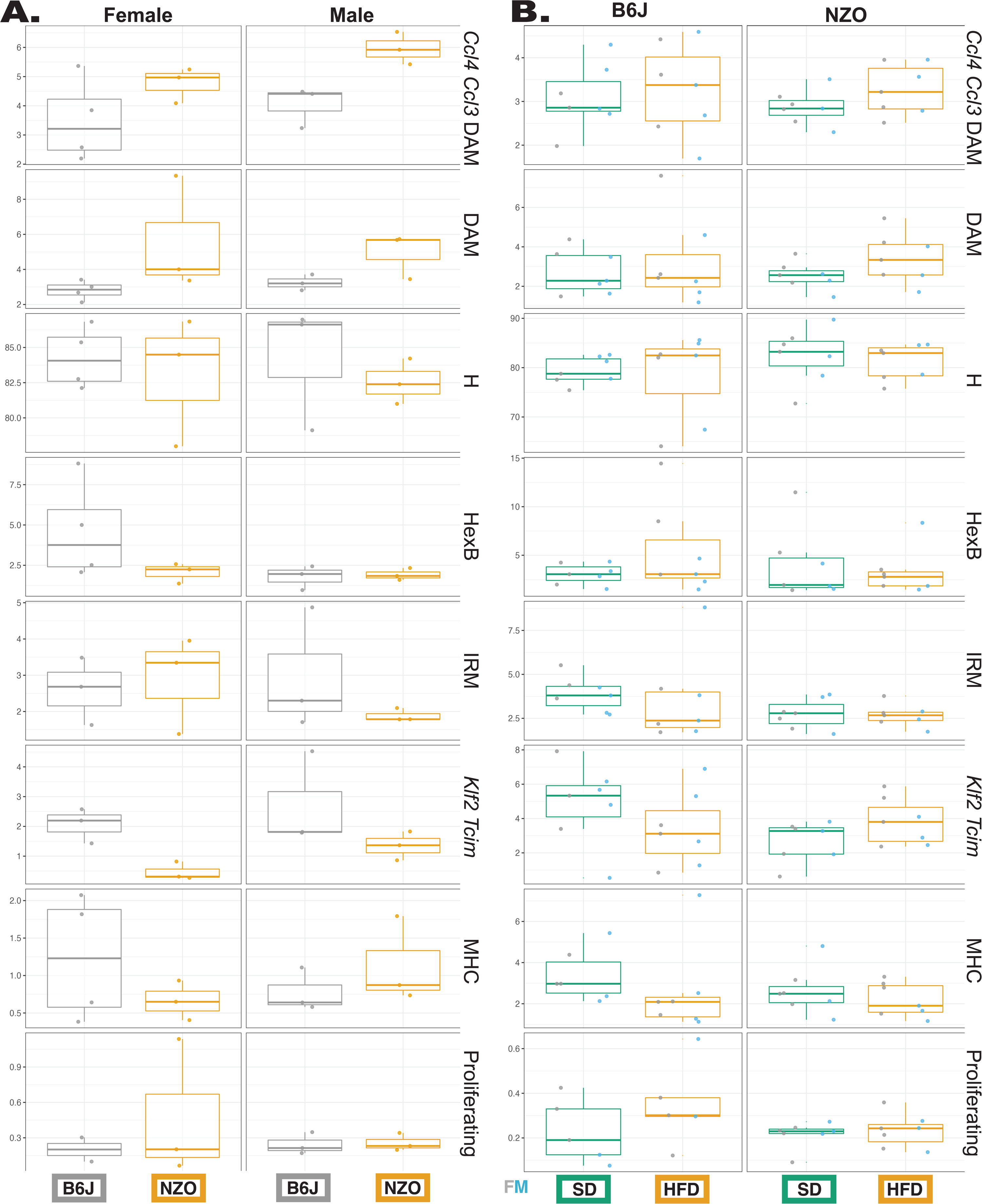

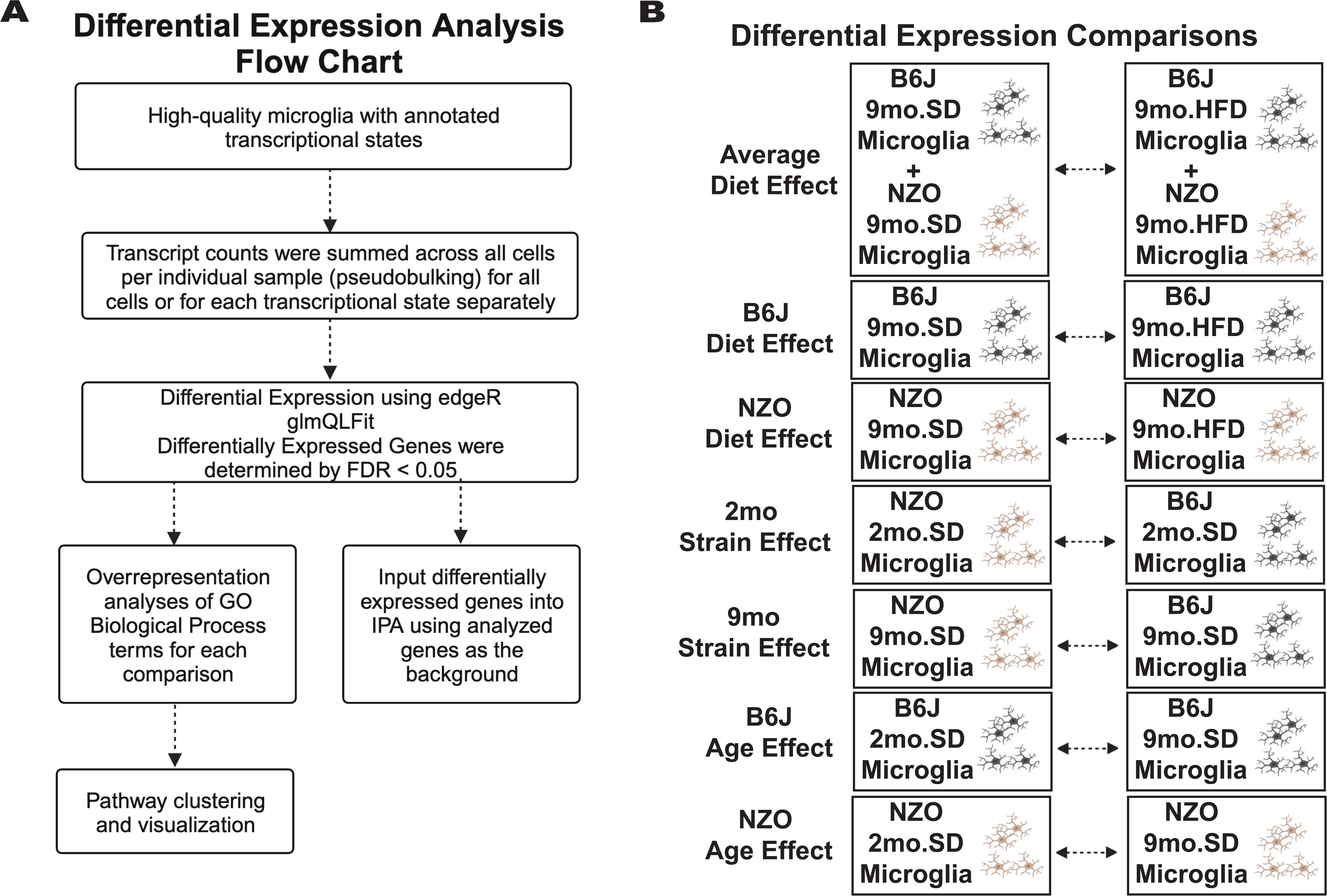

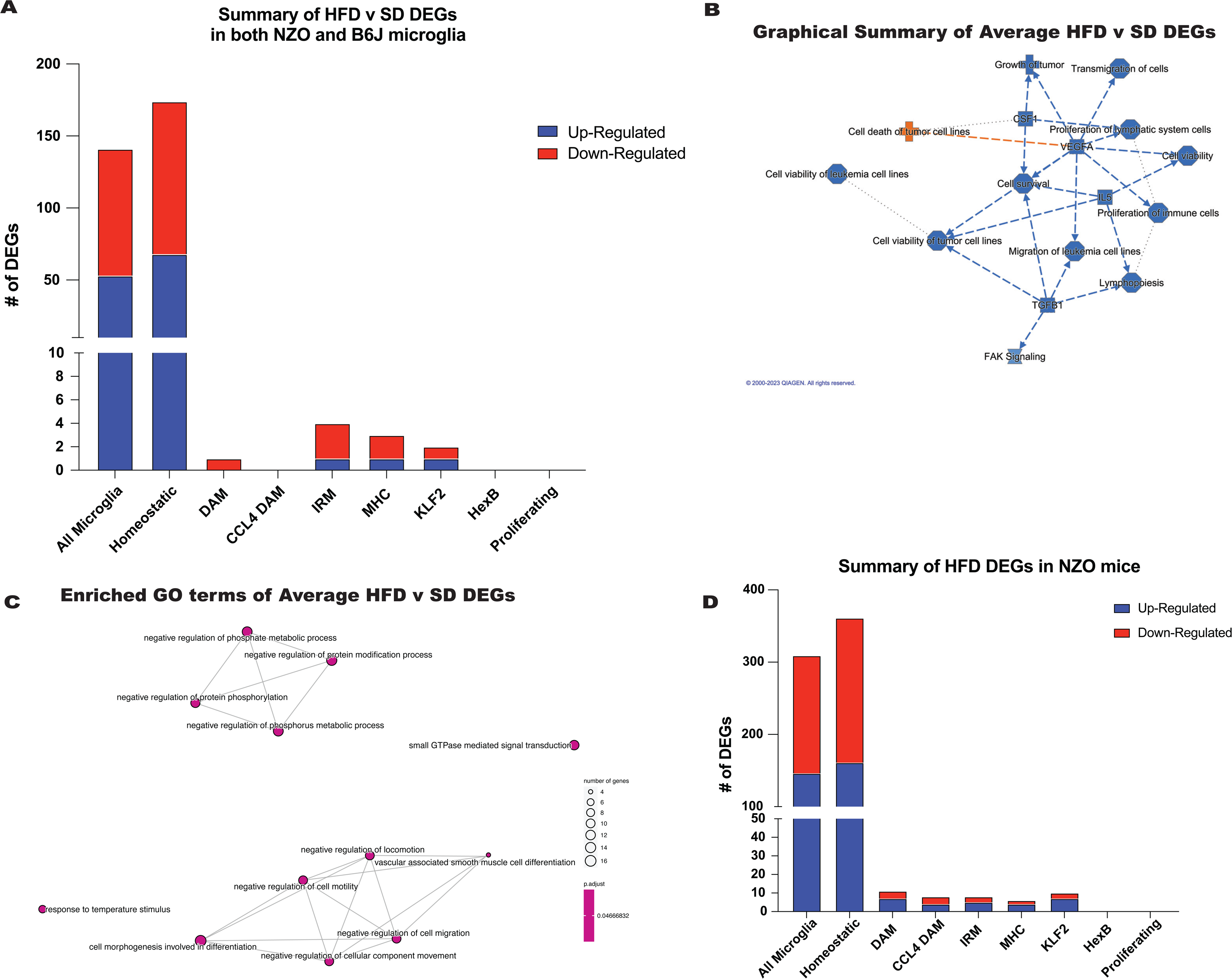

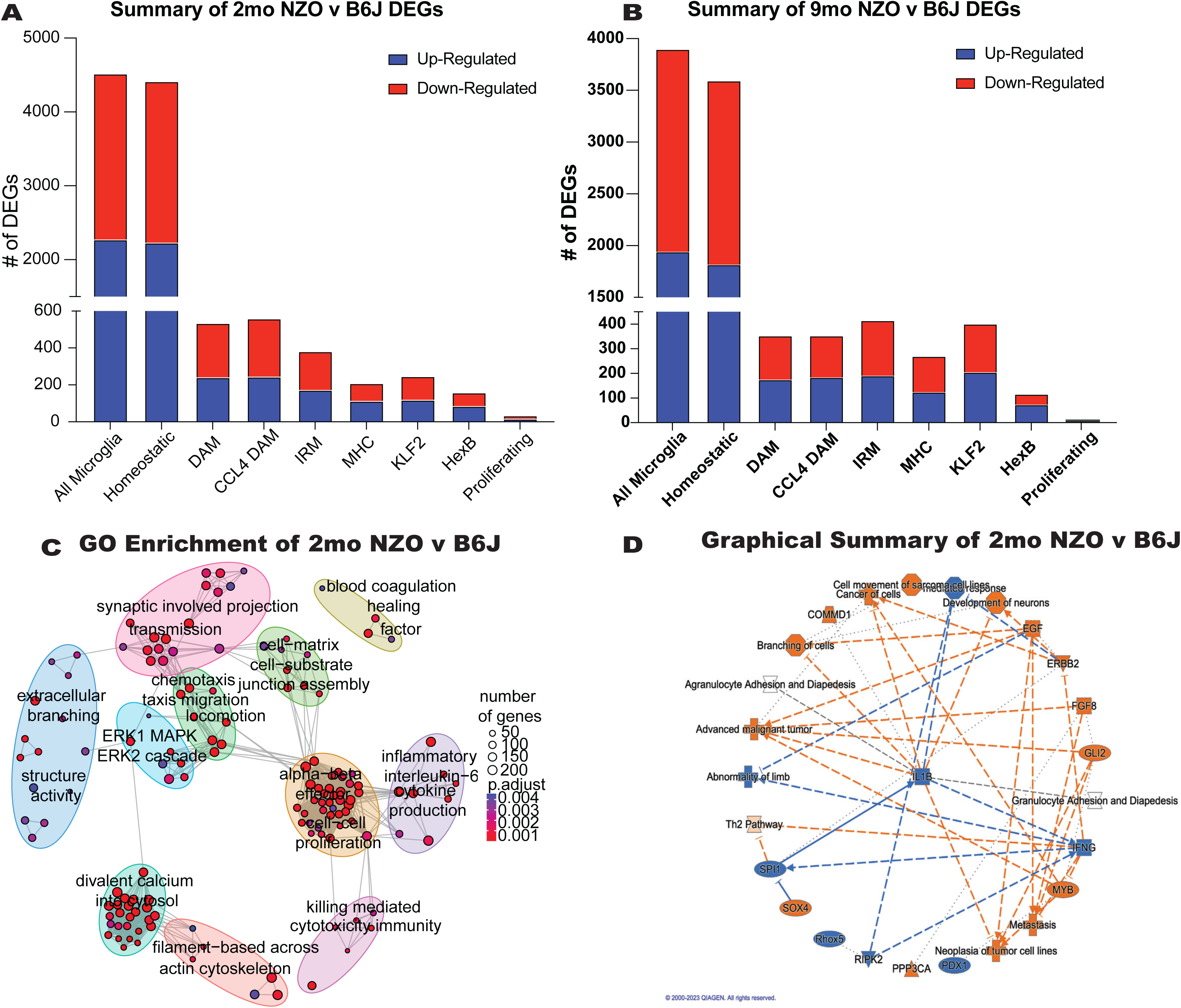

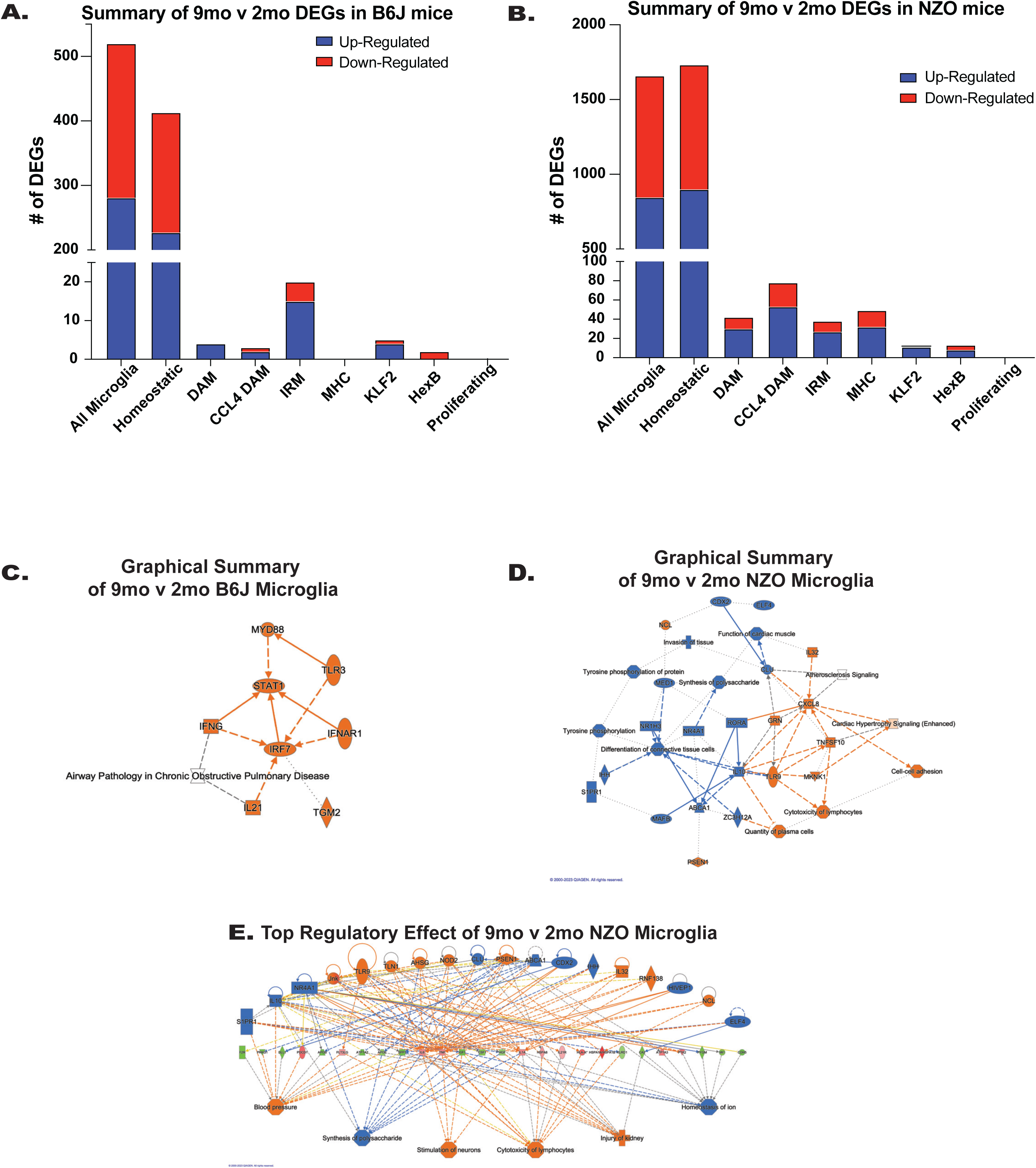

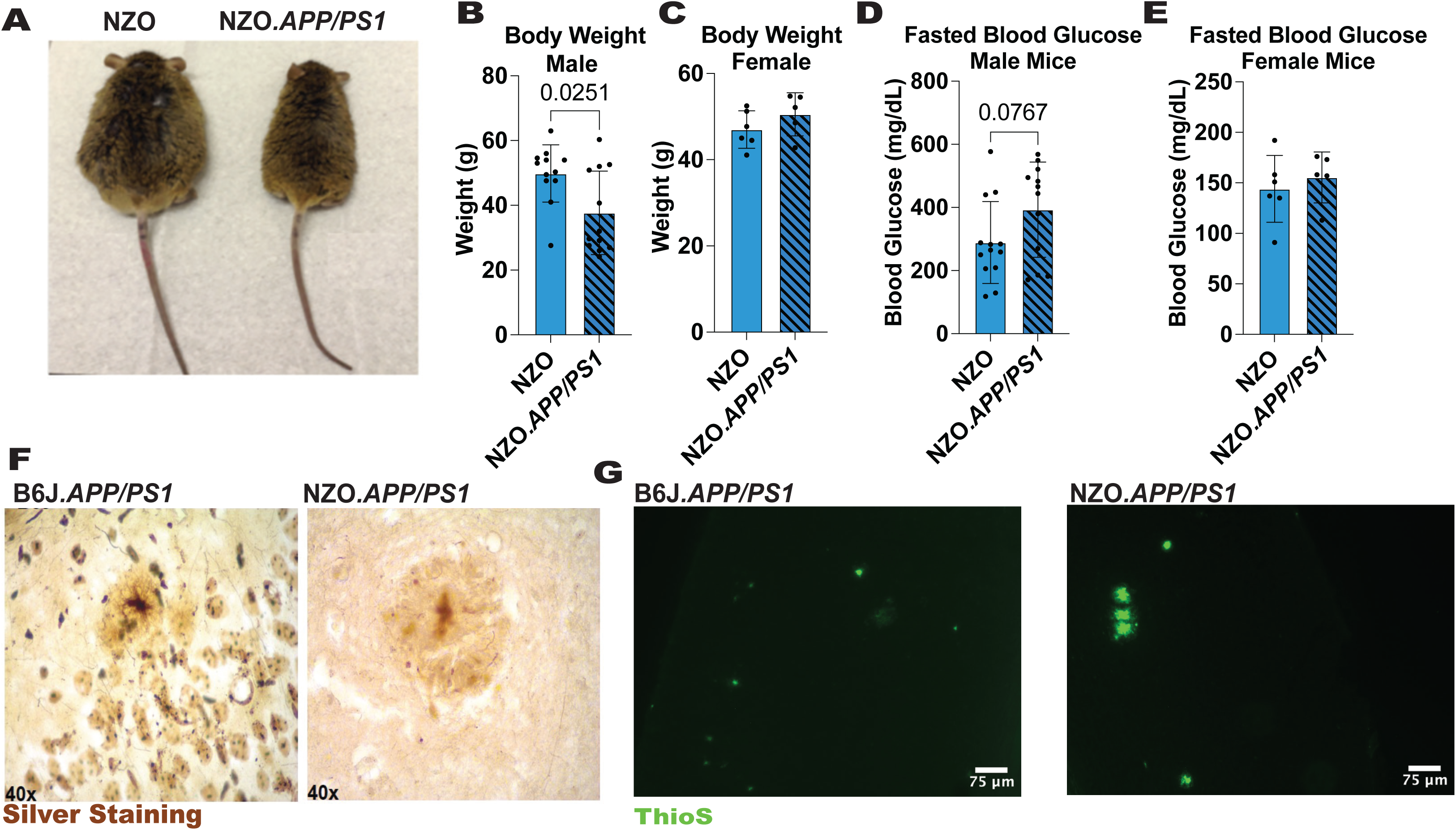

## Notes

### Competing Interest Statement

The authors have declared no competing interest.

