## Supplementary material for "Metabolic syndrome in New Zealand Obese mice promotes microglial-vascular interactions and reduces microglial plasticity": Methods

Resource availability

Lead contact

Further information and requests for resources and reagents should be directed to and will be fulfilled by lead contacts, Gareth Howell or Kristen Onos.

Materials availability

All mouse strains are available through The Jackson Laboratory. All reagents in this study are commercially available.

Data and code availability

Bulk RNA-seq and single cell data are being deposited at GEO. Accession numbers are listed in the key resources table. All the code to replicate the data analysis is available at Zenodo (https://zenodo.org/record/8406521). A shiny app for querying gene expression data from the bulk RNA-seq of brain tissue from NZO and B6J mice treated with LPS or PBS and from the single cell microglia dataset is available at (https://thejacksonlaboratory.shinyapps.io/Howell/).

Experimental model and subject details

Ethics statement

All research was approved by the [Institutional Animal Care and Use Committee](https://www.sciencedirect.com/topics/agricultural-and-biological-sciences/institutional-animal-care-and-use-committee) (IACUC) at The Jackson Laboratory. Authors performed their work following guidelines established by the “The Eighth Edition of the Guide for the Care and Use of Laboratory Animals” and euthanasia using methods approved by the American Veterinary Medical Association.

Mouse strains and cohort generation

All mice were bred and housed in a 12-hour light/dark cycle and received a standard 6% LabDiet Chow (Cat# 5K52) and water ad libitum. A subset of mice was fed a 45% fat diet (ResearchDiets, Cat# D12451i) starting at the age of 2months. Experiments were performed on four mouse strains: C57BL/6J (JAX Stock# 000664), NZO/HILtJ (JAX Stock# 002105), B6.Cg-Tg(APPswe, PSEN1dE9)85Dbo/Mmjax (JAX stock# 005864), and NZO.*APP/PS1* mice generated by backcrossing the Tg(APPswe, PSEN1dE9) transgene to the NZO genetic background for at least 6 generations (1). NZO breeder and NZO.*APP/PS1* mice were fed a 4% LabDiet (Cat# 5K54) chow.

Method details

Metabolic Profiling

For lipid and fasted blood glucose measurements, mice were fasted for 4-6h prior to blood collection. Fasted blood glucose measurements were made using a Contour Next blood glucose monitor (Bayer) using Contour Next Glucose Strips (Bayer, Cat#7278). Blood was collected into to microcentrifuges tubes and allowed to sit at room temperature for 30minutes. Blood was then centrifuged at 2000 x g for 15minutes at room temperature and the serum was collected for analysis. For hemoglobin A1c, unfasted whole blood was collected from mice into EDTA (ThermoFisher, Cat#15575020) coated microcentrifuge tubes for analysis. Total cholesterol, triglycerides, and hemoglobin A1c profiling was completed by the JAX Clinical Chemistry Lab via a Beckman Coulter AU680 chemistry analyzer using standardized tests. Blood pressure was monitored using a non-invasive tail cuff as per the manufacturer’s instructions (CODA Monitor, Kent Scientific). Mice were acclimated to the tail cuff for 5minutes prior to recording. The average of five trials was taken per biological replicate and used for statistical analysis.

Lipopolysaccharide treatment and tissue collection

Lipopolysaccharides (LPS) from E. Coli O111:B4 was re-suspended in sterile PBS at 2.5mg/mL (Sigma-Aldrich, Cat#L3012). 4month old female mice from each strain underwent intraperitoneal injections with either LPS at 5mg LPS / kg body weight or an equivalent volume of PBS. Mice were kept 16h before being anesthetized with Ketamine and Xylazine. Mice then underwent cardiac perfusion with PBS and brains were gently removed, hemisected, and then snap frozen. All tissues were at -80C until further processing.

RNA Isolation, Library Preparation and RNA-Sequencing

Dissected hemibrains were homogenized in MR1 buffer (Macherey-Nagel, Cat#744351.125) using a gentleMACS dissociator (Miltenyi Biotec). Then RNA was isolated from tissue using the NucleoMag RNA Kit (Macherey-Nagel, Cat#744350.4) and the KingFisher Flex purification system (ThermoFisher). RNA isolation was performed according to the manufacturer’s protocol. RNA concentration and quality were assessed using the Nanodrop 8000 spectrophotometer (Thermo Scientific) and the RNA 6000 Nano and Pico Assay (Agilent Technologies). Only samples with RINs > 8 were processed further. Libraries were constructed using the KAPA RNA Hyper Prep Kit with RiboErase (Roche Sequencing and Life Science, Cat#8098131702), according to the manufacturer’s protocol. The quality and concentration of the libraries were assessed using the D5000 ScreenTape (Agilent Technologies) and Qubit dsDNA HS Assay (ThermoFisher, Cat#Q32851), respectively, according to the manufacturers’ instructions. Approximately, 60M 150bp paired-end reads were sequenced per sample on an Illumina NovaSeq 6000 using the S4 Reagent Kit v1.5 (Illumina, Cat#20028313) by the Genome Technologies Core at the Jackson Laboratory.

Hemibrain RNA-Sequencing data analysis

Raw FASTQ files were processed using standard quality control practices (2,3). High quality read pairs were aligned to the mouse genome (mm10) using STAR 2.7.9a (4). A custom genome was generated for NZO mice by incorporating REL-1505 variants into the reference genome (4,5). Read pair counts were summed with the featureCounts function in subread 1.6.3 using the Ensembl Release 68 transcriptome reference (6,7). Differential expression analyses were completed using edgeR v3.40.2 within the R environment v4.2.3 using the glmQLFtest function (8). Groups were defined as a combination of strain and treatment (~0+Group). Contrasts were defined to test for the average treatment effect, the treatment effect within each strain, the average strain effect, and the interaction effect between strain and treatment. Genes were filtered using filterByExpr within edgeR prior to normalization. The clusterProfiler v4.6.2 package within R was used to test for overrepresentation of GO Biological Process gene sets within the list of differentially expressed genes (DEGs) with a false-discovery rate of less than 0.05 and enrichPlot v1.18.4 and EnhancedVolcano v1.16.0 was used to visualize the enrichment results (9-15).

Single myeloid cell preparation

We isolated CD11B+ cells from whole mouse brains using a modified protocol from our previous work (16). All procedures were performed on ice or at 4°C to prevent *ex vivo* activation of myeloid cells (17). Rapid cervical dislocation was performed on mice, then brain tissue was dissected and kept on ice in homogenization buffer: Hanks balanced salt solution (HBSS) (ThermoFisher, Cat#14185-052) containing 15mM HEPES (ThermoFisher, Cat#15630080) and 0.5% glucose (Sigma-Aldrich, Cat#49163) containing 320KU/mL DNaseI (Worthington, Cat#DPRFS). Each brain was minced with a razor blade for initial tissue disruption. Brain tissue from one sample of each strain was pooled together in a 15mL tissue homogenizer in 2mL of homogenization buffer. The tissue was homogenized gently using the loose pestle with 4-5 strokes. The cell suspension was transferred to a new conical tube and passed through a pre-wet 70µm cell strainer (Miltenyi Biotec, Cat#130-110-916). The filtered cell suspension was spun down at 500 x g for 5 minutes at 4°C. The resulting suspension was subjected to debris removal using a debris removal solution (Miltenyi Biotec, Cat#130-109-398) as per the manufacturer’s guidelines. After debris removal, the resulting cell pellet was resuspended in 200µL of myelin removal beads (Miltenyi Biotec, Cat#130-096-733) and was incubated for 10minutes at 4°C. After the incubation, the volume was increased to 2mL and then centrifuged at 500 x g for 5 minutes at 4°C. The cell pellet was resuspended in PBS+ 3% FBS (Fetal Bovine Serum, Gibco, Cat#A3160402) and were transferred to pre-wet LD columns (Miltenyi Biotec, Cat#130-042-901) attached to an QuadroMACS separator (Miltenyi Biotec, Cat#130-090-976). Flow through was collected and then centrifuged at 500 x g for 5 minutes at 4°C. The cell pellet was then incubated with anti-CD11B microbeads (Miltenyi Biotec, Cat#130-049-601) to enrich for brain myeloid cells as per the manufacturer’s guidelines. Resulting cell suspensions were assessed for viability and counted prior to subjecting to scRNA-seq by the JAX Single Cell Laboratory. Viability of all pools submitted for scRNA-seq was greater than 85%.

Single-cell library preparation and RNA-sequencing

CD11B-positive enriched brain myeloid cells were subjected to single-cell library preparation. For each pool an average of 12,400 cells were washed and resuspended in PBS containing 3% FBS and immediately processed as follows. Single-cell capture, barcoding and library preparation were performed using the 10X Chromium platform (10X Genomics), using a Chromium Next GEM Single Cell 3’ High Throughput kit according to the manufacturer’s protocol (10X Genomics, Cat#1000371). The resulting cDNA and indexed libraries were checked for quality on an Agilent 4200 TapeStation, quantified by KAPA qPCR, and sequenced on an Illumina NovaSeq 6000 S4 flow cell, targeting approximately 6,200 barcoded cells with an average sequencing depth of 50,000 reads per cell. Illumina base call (bcl) files for the samples were converted to FASTQ files using CellRanger bcl2fastq (version 2.20.0.422, Illumina). We generated a subset of the larger strain variation recorded in the VCFs for the mouse genome project using bcftools to generate an NZO-B6 VCF (5,18). We then utilized demuxlet with the NZO-B6 VCF to assign a source strain for each cell using the CellRanger BAM files (19). All cells which were ambiguous in origin, e.g., doublets, were discarded from downstream analyses.

Quality control and integration of single cell RNA-seq data

First, cells with less than 600 detected genes, or higher than 8% mitochondrial genes were removed using Seurat v4.9.9 (20,21). 20 10X batches were merged into one Seurat object, then split by Strain. The data were then integrated together and normalized using the sctransform (SCT) and reduction= “rpca” (22). Uniform manifold approximation and projection (UMAP) dimensionality reduction was performed on the integrated data, and clusters were identified using FindNeighbors with the first 99 principal components and FindClusters with a resolution of 0.8. One cluster of cells originated entirely from one mouse (with 2500 cells from one mouse, and with less than 50 cells from any other mouse). Cells from this mouse were discarded and dimensionality reduction and clustering was re-performed. 28 clusters were identified. Cell types were assigned using marker genes and SingleR v2.0, which compares the whole-transcriptome profiles of each cell to expression profiles of known, purified cell types using the ImmGen database available in the celldex package v1.8 (23,24). We identified putative populations of microglia (83,757), neutrophils (9,464), monocytes (2,379), and other immune cell types (<1,000 total). On putative microglia we performed another round of integration, using SCT normalization and reduction = “cca” to improve integration of the microglia across the strains. Dimensionality reduction, FindNeighbors, and FindClusters were performed as above. Small clusters of border associated macrophages (*Mrc1*+), erythrocytes (*Hbb*+, *Hba*+), and neutrophils (*Ly6g+, Selp+, Cxcr2+)* and non-immune cells (*Ptprc*-) were identified and discarded. The resulting 18 sub-clusters were annotated using known marker genes for previously identified microglial states and with new positive marker genes identified with FindAllMarkers in Seurat using the Wilcoxon Rank Sum test. Violin and UMAP plots were generated using the scCustomize v 1.1.1 R package (25). We tested for differences in the proportion of microglia assigned to each transcriptional state using glmQLFtest within edgeR v3.40.2 with library size normalization; we created a model (~ 0 + group + batch) to test contrasts for the average age, strain, and diet effect (8).

Differential gene expression and ingenuity pathway analysis

The single-cell microglia gene raw counts from a given annotated transcriptional state, or from all microglia, of each sample were summed to generate pseudobulked expression profiles (26). These pseudobulked expression profiles then underwent a standard RNA-seq differential expression analysis within edgeR v3.40.2 using glmQLFtest (8). Groups were defined as a combination of strain, sex, diet, and age and batch was included as a covariate (~0+Group+Batch). Contrasts were defined to test for the average diet effect, diet effects with each strain, the age effect within each strain, and strain differences at each age. Genes were filtered using filterByExpr within edgeR prior to normalization. The clusterProfiler v4.6.2 package within R was used to test for overrepresentation of GO Biological Process gene sets within the list of differentially expressed genes (DEGs) with a false-discovery rate of less than 0.05 and enrichPlot v1.18.4 was used to visualize the enrichment results (9-12,14).

Gene expression results, including FDR and log_2_fold changes, for each comparison was uploaded to the Ingenuity Pathway Analysis (IPA) software suite (Qiagen Inc.) and used for input for all IPA analyses including graphical summary, regulatory effects, and upstream regulators.

Tissue harvesting and sectioning for immunohistochemistry and histology.

Mice were anesthetized with Ketamine and Xylazine and then underwent rapid cardiac perfusion with PBS. Brains were gently removed and fixed in 4% paraformaldehyde (Electron Microscopy Services, Cat#15714) overnight at 4°C. After fixation, the brain tissue underwent a series of sucrose incubations. First, with 15% sucrose in PBS for 24 h followed by incubation with 30% sucrose in PBS for an additional 24 h. Brains were then frozen and stored at -80°C until utilized. 25µm thick sections from frozen brain tissue were cut using a freezing microtome (ThermoScientific, Microm HM430) and stored in a cryoprotectant solution (31.25% glycerol and 31.25% ethylene glycol in PBS) at 4°C until stained.

Immunohistochemistry

Sections were washed 1× with PBS for 5 min and washed 3× with PBS+1% Triton-X100 (PBS-T) for 5min. Blocking was conducted with 5% Donkey Serum (Sigma-Aldrich, Cat#D9663) in PBS-T for 1h at RT. All primary antibodies were diluted in 2.5% Donkey Serum in PBS-T and applied overnight at 4°C. Subsequently, slides were washed 3× with PBS-T, then incubated with secondary antibodies diluted in PBS-T for 1.5h at RT. For slides containing DAPI, sections were washed and placed in a 1 μg/mL DAPI (Invitrogen, Cat#D1306) solution in PBS for 5 min at RT and washed 3× with PBS before mounting sections onto glass slides with fluorescent mounting media (Polysciences, Cat#18606-20).

Primary antibodies used: 1µg/mL Rabbit Anti-IBA1 (Wako, Cat#019-19741), 2µg/mL Chicken Anti-TMEM119 (Synaptic Systems, Cat#400 006), 10µg/mL Rabbit Anti-Fibrin/Fibrinogen (Dako, Cat#A0080), 1:500 (v/v) Mouse Anti-β-Amyloid (Clone 6E10, Biolegend, Cat#803015), 2µg/mL Goat anti-CD31 (R&D Systems, Cat#AF3628), 4µg/mL Rat anti-LAMP1 (Abcam, Cat#ab25245). Secondary antibodies utilized: Donkey anti-Mouse Alexa Fluor 488 (Invitrogen, Cat#A21203), Donkey anti-Rat Alexa Fluor 488 (Invitrogen, Cat#A-21208), Donkey anti-Goat Alexa Fluor 488 (Abcam, Cat#ab150133), Donkey anti-Mouse Alexa Fluor 568 (Invitrogen, Cat#A10037), Donkey anti-Rabbit Alexa Fluor 568 (Invitrogen, Cat#A10042), Donkey anti-Rabbit Alexa Fluor 647 (Invitrogen, Cat#A-31573), Donkey anti-Chicken Alexa Fluor 647 (Jackson ImmunoResearch, Cat#703-605-155). All secondary antibodies were used at 1 μg/mL. All experiments included negative controls by omission of primary antibody.

For Thioflavin S staining, sections were first incubated with 1% Thioflavin S (Sigma-Aldrich, Cat#T1892) dissolved in an equal parts water: ethanol solution for eight minutes at RT, followed 3x washes with 80% ethanol, then 1 wash with 95% ethanol, and a final wash in dH2O, and mounted with fluorescent mounting medium.

Imaging and quantification

Multicolor wide-field images were taken on a Leica DMi8 microscope at × 20 or × 40 magnification. Thioflavin S images were taken on a Zeiss Axio Imager Z2 at × 10 magnification. High resolution images were taken on a Leica SP8 confocal microscope at × 63 magnification. All microscope settings were kept identical for each experiment. 2–4 images/tissue slice and 3–4 tissue slices/sample were collected for analysis. All analyses were performed with the Fiji distribution of ImageJ (NIH)(27,28).

Area and Counts analyses

To quantify the positive area of each stain, the images underwent background removal with the rolling ball radius set to 50 within ImageJ. Images were then subjected to automated thresholding with the optimal thresholding algorithm of each signal being selected by a naïve experimenter. The following thresholding algorithms were utilized to determine positive area: Triangle used for IBA1, TMEM119, and CD31, Yen for Fibrin/Fibrinogen, and Otsu used for DAPI and 6E10 (29-31). Calculation of positive area was calculated as percent of the field of view. Automated TMEM119^+^ DAPI^+^ cell counts and 6E10^+^ amyloid plaques were completed by counting the number of overlapping positive objects using the Analyze Particles function within Fiji with minimum size cutoffs of 5µm^2^ or 10µm^2^, respectively. The average of 4-6 images per unique biological sample was used for statistical analysis.

Histology

Sections were mounted onto glass slides and dried at RT overnight. Slides were stained with Prussian Blue using the Iron Stain Kit (Abcam, Cat#ab150674) as per the manufacturer’s guidelines. Stained slides were dehydrated with 3 washes with reagent alcohol and then mounted using organic mounting medium (Organo/Limonene Mount, Sigma-Aldrich, Cat#O8015). Images were taken using a Nikon Eclipse E200 microscope equipped with a SPOT 5.0 MP camera (Model# 28.2-5MP, Diagnostic Instruments, Inc.). Prussian blue positive spots were manually counted across 4-8 sections per biological sample. The average number of spots per section for each biological replicate were calculated and used for statistical analyses.

Quantification and statistical analysis

Data are shown as mean ± SEM. Normality of the data was assessed using the Shapiro-Wilk’s normality test. For two group comparisons, normal data were subsequently evaluated by independent two-sample *t*-tests; otherwise, data were evaluated by a non-parametric Mann–Whitney test. For longitudinal body weight data, a mixed effects model was used to test for strain, sex, and diet effects. For blood metabolite and blood pressure data, a two- or three-way ANOVA was performed as indicated to examine strain, sex, and in some cases, diet, and their interaction effects. All analyses were performed with GraphPad Prism 9 (GraphPad Software).

1.	Jankowsky, J. L., Fadale, D. J., Anderson, J., Xu, G. M., Gonzales, V., Jenkins, N. A., Copeland, N. G., Lee, M. K., Younkin, L. H., Wagner, S. L., Younkin, S. G., and Borchelt, D. R. (2004) Mutant presenilins specifically elevate the levels of the 42 residue beta-amyloid peptide in vivo: evidence for augmentation of a 42-specific gamma secretase. *Hum Mol Genet* **13**, 159-170

2.	Andrews, S. (2010) FastQC: a quality control tool for high throughput sequence data.

3.	Wingett, S. W., and Andrews, S. (2018) FastQ Screen: A tool for multi-genome mapping and quality control. *F1000Res* **7**, 1338

4.	Dobin, A., Davis, C. A., Schlesinger, F., Drenkow, J., Zaleski, C., Jha, S., Batut, P., Chaisson, M., and Gingeras, T. R. (2013) STAR: ultrafast universal RNA-seq aligner. *Bioinformatics* **29**, 15-21

5.	Keane, T. M., Goodstadt, L., Danecek, P., White, M. A., Wong, K., Yalcin, B., Heger, A., Agam, A., Slater, G., Goodson, M., Furlotte, N. A., Eskin, E., Nellaker, C., Whitley, H., Cleak, J., Janowitz, D., Hernandez-Pliego, P., Edwards, A., Belgard, T. G., Oliver, P. L., McIntyre, R. E., Bhomra, A., Nicod, J., Gan, X., Yuan, W., van der Weyden, L., Steward, C. A., Bala, S., Stalker, J., Mott, R., Durbin, R., Jackson, I. J., Czechanski, A., Guerra-Assuncao, J. A., Donahue, L. R., Reinholdt, L. G., Payseur, B. A., Ponting, C. P., Birney, E., Flint, J., and Adams, D. J. (2011) Mouse genomic variation and its effect on phenotypes and gene regulation. *Nature* **477**, 289-294

6.	Cunningham, F., Allen, J. E., Allen, J., Alvarez-Jarreta, J., Amode, M. R., Armean, I. M., Austine-Orimoloye, O., Azov, A. G., Barnes, I., Bennett, R., Berry, A., Bhai, J., Bignell, A., Billis, K., Boddu, S., Brooks, L., Charkhchi, M., Cummins, C., Da Rin Fioretto, L., Davidson, C., Dodiya, K., Donaldson, S., El Houdaigui, B., El Naboulsi, T., Fatima, R., Giron, C. G., Genez, T., Martinez, J. G., Guijarro-Clarke, C., Gymer, A., Hardy, M., Hollis, Z., Hourlier, T., Hunt, T., Juettemann, T., Kaikala, V., Kay, M., Lavidas, I., Le, T., Lemos, D., Marugan, J. C., Mohanan, S., Mushtaq, A., Naven, M., Ogeh, D. N., Parker, A., Parton, A., Perry, M., Pilizota, I., Prosovetskaia, I., Sakthivel, M. P., Salam, A. I. A., Schmitt, B. M., Schuilenburg, H., Sheppard, D., Perez-Silva, J. G., Stark, W., Steed, E., Sutinen, K., Sukumaran, R., Sumathipala, D., Suner, M. M., Szpak, M., Thormann, A., Tricomi, F. F., Urbina-Gomez, D., Veidenberg, A., Walsh, T. A., Walts, B., Willhoft, N., Winterbottom, A., Wass, E., Chakiachvili, M., Flint, B., Frankish, A., Giorgetti, S., Haggerty, L., Hunt, S. E., GR, I. I., Loveland, J. E., Martin, F. J., Moore, B., Mudge, J. M., Muffato, M., Perry, E., Ruffier, M., Tate, J., Thybert, D., Trevanion, S. J., Dyer, S., Harrison, P. W., Howe, K. L., Yates, A. D., Zerbino, D. R., and Flicek, P. (2022) Ensembl 2022. *Nucleic Acids Res* **50**, D988-D995

7.	Liao, Y., Smyth, G. K., and Shi, W. (2013) The Subread aligner: fast, accurate and scalable read mapping by seed-and-vote. *Nucleic Acids Res* **41**, e108

8.	Robinson, M. D., McCarthy, D. J., and Smyth, G. K. (2010) edgeR: a Bioconductor package for differential expression analysis of digital gene expression data. *Bioinformatics* **26**, 139-140

9.	Ashburner, M., Ball, C. A., Blake, J. A., Botstein, D., Butler, H., Cherry, J. M., Davis, A. P., Dolinski, K., Dwight, S. S., Eppig, J. T., Harris, M. A., Hill, D. P., Issel-Tarver, L., Kasarskis, A., Lewis, S., Matese, J. C., Richardson, J. E., Ringwald, M., Rubin, G. M., and Sherlock, G. (2000) Gene ontology: tool for the unification of biology. The Gene Ontology Consortium. *Nat Genet* **25**, 25-29

10.	Gene Ontology, C., Aleksander, S. A., Balhoff, J., Carbon, S., Cherry, J. M., Drabkin, H. J., Ebert, D., Feuermann, M., Gaudet, P., Harris, N. L., Hill, D. P., Lee, R., Mi, H., Moxon, S., Mungall, C. J., Muruganugan, A., Mushayahama, T., Sternberg, P. W., Thomas, P. D., Van Auken, K., Ramsey, J., Siegele, D. A., Chisholm, R. L., Fey, P., Aspromonte, M. C., Nugnes, M. V., Quaglia, F., Tosatto, S., Giglio, M., Nadendla, S., Antonazzo, G., Attrill, H., Dos Santos, G., Marygold, S., Strelets, V., Tabone, C. J., Thurmond, J., Zhou, P., Ahmed, S. H., Asanitthong, P., Luna Buitrago, D., Erdol, M. N., Gage, M. C., Ali Kadhum, M., Li, K. Y. C., Long, M., Michalak, A., Pesala, A., Pritazahra, A., Saverimuttu, S. C. C., Su, R., Thurlow, K. E., Lovering, R. C., Logie, C., Oliferenko, S., Blake, J., Christie, K., Corbani, L., Dolan, M. E., Drabkin, H. J., Hill, D. P., Ni, L., Sitnikov, D., Smith, C., Cuzick, A., Seager, J., Cooper, L., Elser, J., Jaiswal, P., Gupta, P., Jaiswal, P., Naithani, S., Lera-Ramirez, M., Rutherford, K., Wood, V., De Pons, J. L., Dwinell, M. R., Hayman, G. T., Kaldunski, M. L., Kwitek, A. E., Laulederkind, S. J. F., Tutaj, M. A., Vedi, M., Wang, S. J., D'Eustachio, P., Aimo, L., Axelsen, K., Bridge, A., Hyka-Nouspikel, N., Morgat, A., Aleksander, S. A., Cherry, J. M., Engel, S. R., Karra, K., Miyasato, S. R., Nash, R. S., Skrzypek, M. S., Weng, S., Wong, E. D., Bakker, E., Berardini, T. Z., Reiser, L., Auchincloss, A., Axelsen, K., Argoud-Puy, G., Blatter, M. C., Boutet, E., Breuza, L., Bridge, A., Casals-Casas, C., Coudert, E., Estreicher, A., Livia Famiglietti, M., Feuermann, M., Gos, A., Gruaz-Gumowski, N., Hulo, C., Hyka-Nouspikel, N., Jungo, F., Le Mercier, P., Lieberherr, D., Masson, P., Morgat, A., Pedruzzi, I., Pourcel, L., Poux, S., Rivoire, C., Sundaram, S., Bateman, A., Bowler-Barnett, E., Bye, A. J. H., Denny, P., Ignatchenko, A., Ishtiaq, R., Lock, A., Lussi, Y., Magrane, M., Martin, M. J., Orchard, S., Raposo, P., Speretta, E., Tyagi, N., Warner, K., Zaru, R., Diehl, A. D., Lee, R., Chan, J., Diamantakis, S., Raciti, D., Zarowiecki, M., Fisher, M., James-Zorn, C., Ponferrada, V., Zorn, A., Ramachandran, S., Ruzicka, L., and Westerfield, M. (2023) The Gene Ontology knowledgebase in 2023. *Genetics* **224**

11.	Wu, T., Hu, E., Xu, S., Chen, M., Guo, P., Dai, Z., Feng, T., Zhou, L., Tang, W., Zhan, L., Fu, X., Liu, S., Bo, X., and Yu, G. (2021) clusterProfiler 4.0: A universal enrichment tool for interpreting omics data. *Innovation (Camb)* **2**, 100141

12.	Yu, G., Wang, L. G., Han, Y., and He, Q. Y. (2012) clusterProfiler: an R package for comparing biological themes among gene clusters. *OMICS* **16**, 284-287

13.	Benjamini, Y., and Hochberg, Y. (1995) Controlling the False Discovery Rate: A Practical and Powerful Approach to Multiple Testing. *Journal of the Royal Statistical Society. Series B (Methodological)* **57**, 289-300

14.	Yu, G. (2023) enrichplot: Visualization of Functional Enrichment Result.

15.	Blighe K, R. S., Lewis M. (2023) EnhancedVolcano: Publication-ready volcano plots with enhanced colouring and labeling.

25.	Marsh, S. E. (2021) scCustomize: Custom Visualizations & Functions for Streamlined Analyses of Single Cell Sequencing.
