## Supplemental Figure Legends for "Metabolic syndrome in New Zealand Obese mice promotes microglial-vascular interactions and reduces microglial plasticity"

*Figure S1. Single cell RNA-sequencing from NZO and B6J central nervous system myeloid cells.*

**a.** Dimensionality reduction plot (UMAP) of sequenced cells colored by cluster identity after integration and application of quality control metrics. **b.** Violin plots showing number of genes, RNA counts, and percent mitochondrial reads. **c.** Dot plot of marker gene expression for microglia, monocytes, macrophages, granulocytes, erythrocytes, NK cells, and B cells. **d.** UMAP of sequenced cells colored by predicted cell-type determined by SingleR using the Immgen database as a reference. **e.** UMAP of re-clustered microglia colored by assigned cluster. **f.** Dot plot of marker gene expression for each cluster. 9mo mice: *N*=6M NZO (3SD,3HFD), *N*=8F NZO (4SD,4HFD), *N*=8M B6J (4SD,4HFD), *N*=6F B6J (3SD,3HFD); 2mo mice: *N*=6 NZO mice (3F,3M), *N=*7 B6J mice (4M,3F).

*Figure S2. Proportions of microglia assigned to each transcriptional state across each age, sex, strain, and diet.*

**a.** Boxplots illustrating proportion of assigned microglia to each microglial transcriptional state out of total microglia captured at 2mo with the plots split by sex and colored by strain. *N*=6 NZO (3F,3M), *N=*7 B6J mice (4M,3F). **b.** Boxplots illustrating proportion of assigned microglia to each microglial transcriptional state out of total microglia captured at 9mo with the plots split by strain and colored by diet. *N*=6M NZO (3SD,3HFD), *N*=8F NZO (4SD,4HFD), *N*=8M B6J (4SD,4HFD), *N*=6F B6J (3 SD,3HFD) mice.

*Figure S3. Illustrations of differential expression workflow.*

**a.** Flow chart illustrating differential expression and secondary analyses. **b.** Illustrations of the various comparisons made for differential expression analyses. For **a-b**, Created with BioRender.com.

*Figure S4. The average HFD effect across strains is associated with modest changes in differential gene expression profiles and is associated with cell survival and cellular movement pathways.*

**a.** Bar chart summarizing the number of DEGs associated with the HFDvSD in both strains across all microglia or within each individual microglia annotated state. **b.** IPA graphical summary of HFDvSD DEGs. **c.** Enrichment plot of GO terms from *a* for all microglia. **d.** Bar chart summarizing the number of DEGs associated with HFDvSD across all microglia or within each individual microglia annotated state in only NZO mice. *N*=14SD (3M NZO, 4F NZO, 4M B6J, 3F B6J) and *N*=14HFD (3M NZO, 4F NZO, 4M B6J, 3F B6J) mice.

*Figure S5. NZO and B6J microglia exhibit large strain-associated transcriptional differences.*

Bar charts summarizing the number of DEGs associated with NZOvB6J across all microglia or within each individual microglia annotated state within SD fed animals at (**a**) 2mo or (**b**) 9mo. **c.** Enrichment GO term plot from 2mo NZOvB6J DEGs for all microglia. Colored circles indicate pathway cluster identity. Key words for each pathway cluster are displayed. **d.** IPA graphical summary of 2mo NZOvB6J DEGs for all microglia. 9mo mice: N=7 NZO (3M,4F), N=7 B6J (4M,3F). 2mo mice: N=6 NZO mice (3M,3F), N=7 (3M,4F) B6J mice.

*Figure S6. Aging from 2-9months influences microglia in a strain dependent manner.*

Bar charts summarizing the number of DEGs associated with 9v2mo microglia across all microglia or within each individual microglia annotated state for SD fed B6J mice (**a**) or NZO (**b**). IPA graphical summaries of 9v2mo DEGs in all B6J (**c**) or NZO (**d**) microglia. **e**. Top IPA regulatory effect for the 9v2mo NZO all microglia DEGs. 9mo mice: N=7 NZO (3M,4F), N=7 B6J (4M,3F). 2mo mice: N=6 NZO mice (3/sex), N=7 (3M,4F) B6J mice.

*Figure S7. Male NZO.APP/PS1 animals display a marked reduction in body weight and exhibit larger and more diffuse plaques than male B6J.APP/PS1animals.*

**a.** Representative images of an NZO and an NZO*.APP/PS1* male mouse. Quantification of body weight of NZO and NZO*.APP/PS1*mice at 6mo in males (**b**) and females (**c**). Fasted blood glucose measurements in NZO and NZO*.APP/PS1*mice at 6mo in males (**d**) and females (**e**). **f.** Representative images of silver staining of the indicated male mouse groups at 8mo of age at 40x magnification. **g.** Representative images of Thioflavin S (ThioS) staining of the indicated male mouse groups at 8mo of age. In **b**, *N*=12M NZO mice/genotype. In **c** and **e**, *N*=6F NZO and N=6F NZO.*APP/PS1* mice. In **d**, *N*=13M NZO and N=13M NZO.*APP/PS1* mice. Independent two sample two-sided *t* test. In **b**, Mann-Whitney test. Data are presented as Mean*±*SD.
